# FertilityOnline, a straight pipeline for functional gene annotation and disease mutation discovery, identifies novel infertility causative mutations in *SYCE1* and *STAG3*

**DOI:** 10.1101/2020.08.05.238162

**Authors:** Jianing Gao, Huan Zhang, Xiaohua Jiang, Asim Ali, Daren Zhao, Jianqiang Bao, Long Jiang, Furhan Iqbal, Qinghua Shi, Yuanwei Zhang

**Affiliations:** The First Affiliated Hospital of USTC, Hefei National Laboratory for Physical Sciences at the Microscale, The CAS Key Laboratory of Innate Immunity and Chronic Diseases, School of Life Sciences, CAS Center for Excellence in Molecular Cell Science, University of Science and Technology of China, Collaborative Innovation Center of Genetics and Development, Hefei 230027, Anhui, China

**Keywords:** Infertility, Database, Functional gene, Mutation

## Abstract

Exploring the genetic basis of human infertility is currently under intensive investigation. However, only a handful of genes are validated in animal models as disease-causing genes in infertile men. Thus, to better understand the genetic basis of spermatogenesis in human and to bridge the knowledge gap between human and other animal species, we have constructed FertilityOnline database, which is a resource that integrates the functional genes reported in literature related to spermatogenesis into an existing spermatogenic database, SpermatogenesisOnline 1.0. Additional features like functional annotation and statistical analysis of genetic variants of human genes, are also incorporated into FertilityOnline. By searching this database, users can focus on the top candidate genes associated with infertility and can perform enrichment analysis to instantly refine the number of candidates in a user-friendly web interface. Clinical validation of this database is established by the identification of novel causative mutations in *SYCE1* and *STAG3* in azoospermia men. In conclusion, FertilityOnline is not only an integrated resource for analysis of spermatogenic genes, but also a useful tool that facilitates to study underlying genetic basis of male infertility.

**Availability:** FertilityOnline can be freely accessed at http://mcg.ustc.edu.cn/bsc/spermgenes2.0/index.html.

## Introduction

Human infertility affects 10-15% of couples at reproductive age, half of which is attributed to the male partner [1, 2]. Spermatogenesis is a delicate, prolonged cell differentiation process that involves self-renewal of spermatogonial stem cells (SSC), meiosis, and post-meiotic development. Disruption of any step during this period likely results in reduced fertility or complete infertility. For example, defective proliferation of SSC may lead to Sertoli cell only syndrome (SCOS), and genetic interference in spermatocytes can result in spermatocyte development arrest (SDA) [3, 4]. It has been estimated that about 25%-50% cases of male infertility result from genetic abnormalities [5, 6]. A survey of literature revealed that at least 2,000 genes are involved in the process of spermatogenesis [7]. However, to date, only a small number of genetic mutations in men have been validated as bonafide causes of human subfertility/infertility in animal models [8, 9].

With the advent of next generation sequencing (NGS), a multitude of high-throughput methods, such as whole exome sequencing (WES) or whole genome sequencing (WGS), are adopted to search for pathogenic mutations in infertile patients [6, 8–10]. These approaches commonly generate enormous datasets, which requires professional analyses and annotation of bioinformatician. To fulfill this requirement, we have constructed FertilityOnline database, which integrates the functional spermatogenic genes reported in literature into the only existing functional spermatogenic database, SpermatogenesisOnline 1.0 [11]. Apart from the basic annotations for manually curated genes (gene information, protein functional domains, pathway, ortholog and paralog, etc.), new features, such as functional annotation, specific gene expression data in different tissues and testicular cell types, and statistical analyses of genetic variants of human genes, have been incorporated in FertilityOnline. With gene or variant annotation in hand, users can directly filter the annotation list to prioritize the candidate genes of interest associated with infertility and perform in-depth enrichment analysis to refine the number of candidates in a user-friendly Web interface. Thus, FertilityOnline not only serves as an integrated database for functional annotation of genes associated with spermatogenesis, but also provides a solid resource for identification of human disease causing genes.

## Material and Methods

FertilityOnline is a comprehensive and systematic collection of functional annotations of spermatogenesis-related genes from the published literature. Information, such as gene expression, gene mutation, and homologs of spermatogenesis-related genes, are also integrated together into this web resource. The list of data sources used in the construction of this back-end database is provided as Table S1. A visual front-end pipeline has also been developed to facilitate users to put their query and to run analysis (Figure 1).

**Figure 1.**
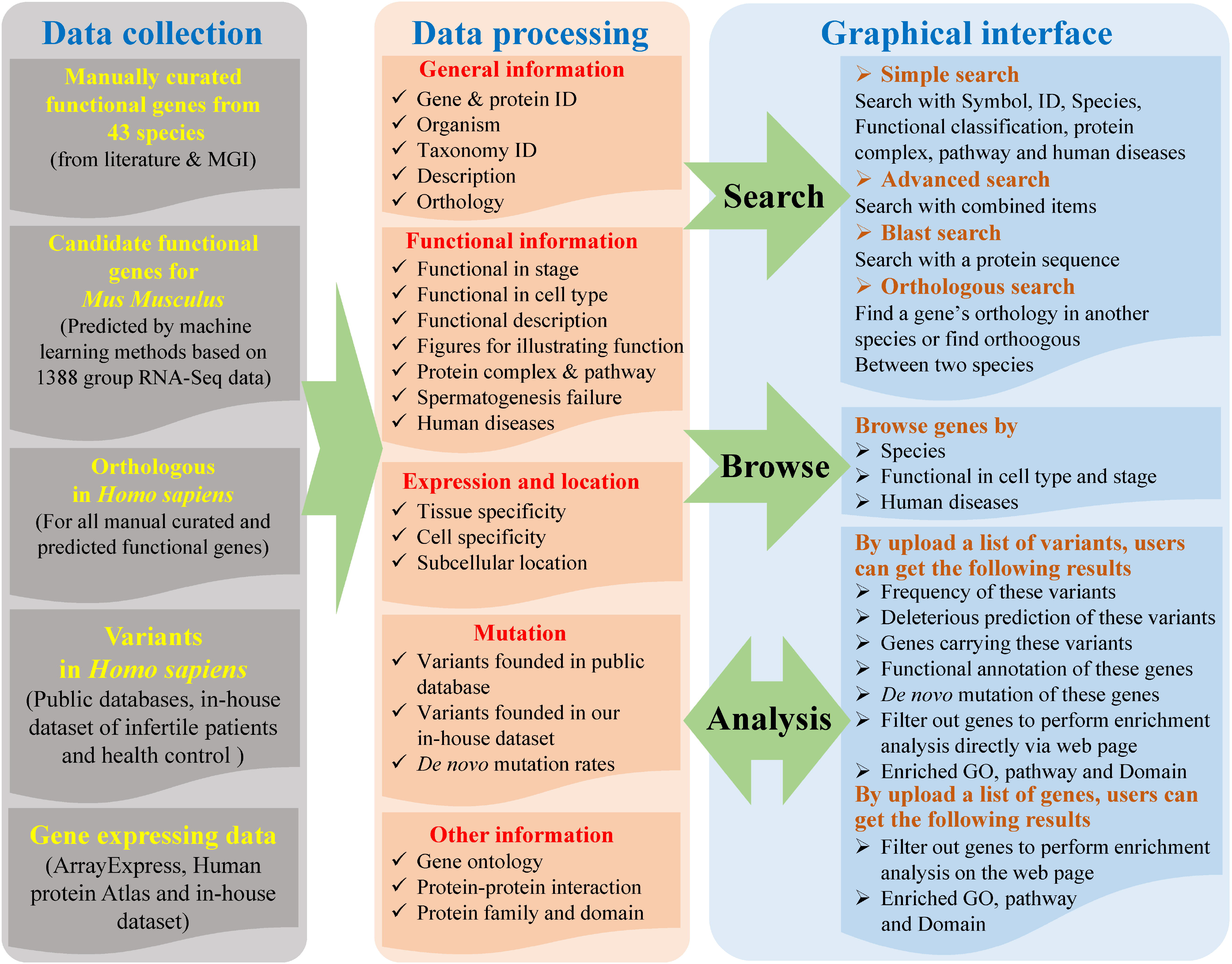
Overall structure of FertilityOnline. FertilityOnline is an integrated database that incorporates information of manually curated functional genes of spermatogenesis and facilitates data processing.

### Data Collection

#### (i) Manually Curated Functional Genes

To comprehensively collect the functional spermatogenic gene information, a number of keywords were employed to search in PubMed database (published before July 1^st^, 2019 in PubMed). For developmental stages, spermatogenesis, spermiogenesis, premeiotic, postmeiotic and meiosis were employed to search the related literature. For cell types in testis, Spermatogonial stem cells (SSC), spermatogonium, spermatogonia, spermatocyte, spermatid, Sertoli cell, Leydig cell and peritubular myoid cell were chosen as keywords. All collected references were manually curated and only the genes with functional experimental validation were deemed as functional genes associated with spermatogenesis. Moreover, figures and tables illustrating the function of these genes were also collected.

#### (ii) Gene Expression Data

The gene expression data collected in this database can be divided into four parts: 1) RNA-Seq data from *Mus musculus* was downloaded from ArrayExpress (Table S2); 2) RNA-Seq data from 37 tissues (appendix, adrenal gland, adipose, bone marrow, colon, cerebral cortex, duodenum, esophagus, gallbladder, heart muscle, kidney, liver, lymph node, lung, ovary, prostate, placenta, pancreas, stomach, spleen, small intestine, skin, salivary gland, thyroid gland, testis, urinary bladder and uterus) of *Homo sapiens* was downloaded from Human Protein Atlas; 3) In-house RNA-Seq data from 5 major mouse testicular cells (spermatogonium, spermatocyte, spermatid, sperm and Sertoli cell); 4) Four sets of public single cell RNA (scRNA)-seq data from human and mouse testes (Table S3). Gene expression data from part 1 was also integrated as features and applied in prediction of candidate functional genes in spermatogenesis (Table S2).

#### (iii) Candidate Functional Genes in Spermatogenesis (Mus musculus)

As mouse is the most widely used model animal in reproductive biology, experimental data accumulated from this species was used for the prediction of candidate functional genes with machine learning method. The positive training dataset contained 653 manually curated genes that were reported to be functional during spermatogenesis. To construct the negative training dataset, we checked the phenotype data from Mouse Genome Informatics (MGI, http://www.informatics.jax.org/), and selected 3,783 genes in which mutation or deletion did not cause any abnormality in reproductive system. The gene expression data (described in ***Gene Expression Data*** part) were used as features to construct the model for predicting candidate functional genes in spermatogenesis. In total, a list of 300 most important features out of 2,627 expression features was employed to train the support vector machine (SVM) model (described in File S1 and Figure S1). Among the predicted positive results, the real positives were defined as true positives (TP), while the others were defined as false positives (FP). As described previously [11], four measurements were adopted to evaluate the performance of our model. The equations are defined below:

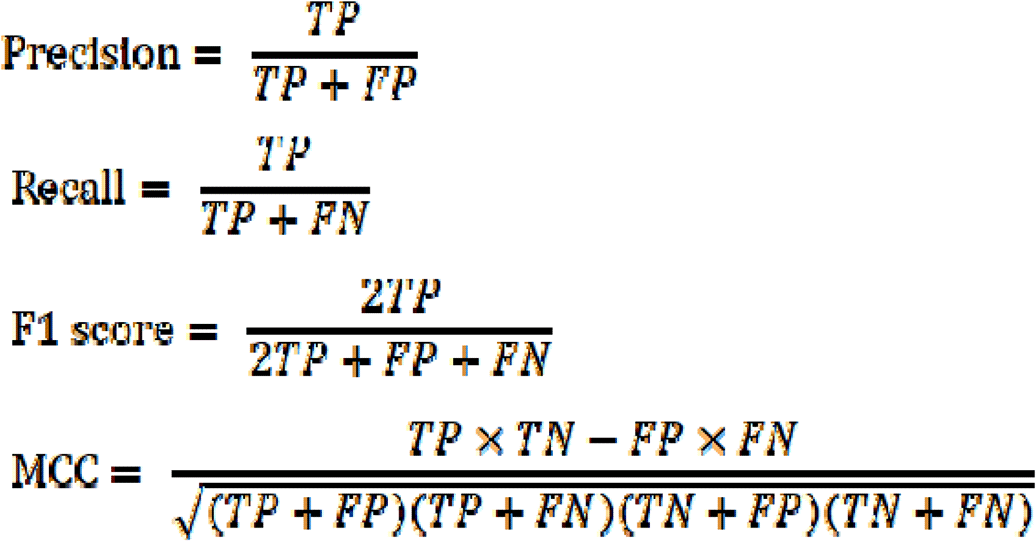

Among the predicted negative results, the real negatives are defined as true negative (TN), while others are defined as false negative (FN). Considering the small training dataset, we perform 4-fold cross-validations rather than 10-fold, and the Receiver Operating Characteristic (ROC) curves were drawn with matplotlib packages.

#### (iv) Orthologous Group Information

Orthologous group information was downloaded from InParanoid (Version 8.0) and PANTHER (Version 12.0) databases. Orthologous groups from these two databases were merged to avoid the loss of group members and redundancy.

#### (v) Variants in Homo sapiens

In FertilityOnline, variants are classified into three categories: 1) variants present in public databases, including 1000G (Phase 3), ExAC (version r0.3.1), ESP6500 (ESP6500SI-V2), UK10K and dbSNP (build 147); 2) variants found in our in-house datasets, including Chinese health control (254 fertile men), European health control (283 fertile Pakistan men), Chinese infertile patients (168 infertile men); 3) background *de novo* mutation rate obtained from Jiang *et al* [12] (Table S1).

### Data Processing

The collected data were processed to provide the following information for each gene:

1. ***General Information***, including gene and protein ID, source organism, taxonomic ID, description and orthology.
2. ***Functional Information***, including functional stage in which it is involve (premeiotic, meiotic and postmeiotic), cell type in which it express (SSC, spermatogonium, spermatocyte, spermatid, Sertoli cell, Leydig cell, etc.), function’s description, figures for illustration of function, protein complex and pathway, spermatogenesis disorder (SCO, SDA and hypospematogenesis (HSG)) and the related human diseases.
3. ***Expression and Localization***, including the normalized value of gene expression in 37 human tissues and their orthologous information in 5 types of mouse testicular cells. We also integrated 4 sets of public scRNA-seq data covering germ and somatic cells. Moreover, the tissue with highest expression is marked and subcellular location information is also provided.
4. ***Mutation***, providing the counts for variants of each gene found in public database as well as our in-house dataset. The *de novo* mutation rates are also provided.
5. ***Other Annotations***, including gene ontology, protein-protein interaction, protein family, domain, etc.

### Implementation

FertilityOnline is hosted on a Dell 730 server, using LAMP architecture (Linux, Apache, MySQL, and PHP). The server is equipped with two 12-core Intel processors (2.2 GHz each) and 128 GB RAM. The backend is supplied by Python and R language and the interface is rendered using jQuery. It takes about 5 minutes to complete an analysis after testing 10 WES generated VCF files (~100,000-400,000 varaints) (Table S4). Additionally, the queuing module can execute more jobs in parallel.

### Exome Sequencing and Data Analysis

Whole Exome Sequencing (WES) was performed on the genomic DNAs (gDNAs) isolated from peripheral blood of non-obstructive azoospermia (NOA) patients using the QIAamp DNA Blood Mini Kit (51206; Qiagen, Hilden, Germany) following the manufacturer’s instructions. An Agilent SureSelect Human All Exon v5 Kit (5190-6208; Santa Clara, CA, USA) was applied to capture the known exons and exon-intron boundary sequences. Sequencing was performed on a Hiseq 2000 platform (Illumina,San Diego, CA, USA) and raw reads (*.fastq format) were aligned to the human reference genome (GRCh37/hg19) using Burrows-Wheeler Aligner (BWA) software by applying default parameters settings. SAM file of each sample was converted to a BAM file by using SAMtools (http://samtools.sourceforge.net/). To remove PCR duplicates and to keep only properly paired reads, Picard tool (http://picard.sourceforge.net/) was used. The Genome Analysis Toolkit (GATK) from Broad Institute (http://www.broadinstitute.org/gatk/) were used to further process the files, and then all BAM files were locally realigned by indel realigner. GATK’s Unified Genotyper was used on the processed BAM files to call both small (INDELs) and single-nucleotide variants (SNVs) within the captured coding exonic intervals. The exome sequencing data has been deposited in ArrayExpress with the accession number of E-MTAB-9287 (described in File S1).

### Western Blotting

To obtain cell lysates, Vero cells were transfected with EGFP-STAG3-WT or EGFP-STAG3-mutant, respectively. Thirty-six hours later, the cells were lyzed and proteins were separated on SDS polyacrylamide gel by electrophoresis for Western blotting as described previously [13].

## Results

### FertilityOnline Integrates Information of Functional Genes in Spermatogenesis

One of the aims of FertilityOnline is to provide an integrated resource that allows users to easily access information about spermatogenic genes and their mutations. To achieve this goal, we collected all the spermatogenic genes reported in the literature by employing a series of keywords to query in PubMed (described in **Materials and Methods**). About 48,000 research articles published before July 1^st^, 2019 were collected. Among them, 4,736 records satisfy the criterion that the function of genes in spermatogenesis is validated by experiment were finally collected in our database. In total, 1610 unique spermatogenic genes with experimental validation from 43 species were curated in our updated database. We found that the functional genes currently reported in spermatogenesis are mainly derived from mouse, which accounts for 61.59% of reported genes, followed by human (15.82%) and rat (10.07%). In contrast, other species comprise the rest of 12.61% altogether (Table S5).

In order to further expand the utilization of FertilityOnline, a prediction model was constructed to infer the candidate functional spermatogenic genes. In this model, functional genes reported in mice were used as positive records, and the genes without any reproductive phenotype after knockout experiments were used as negative records (recorded in the MGI database), and the expression of these genes in 2,627 RNA-Seq datasets (described in **File S1**) were used as features (the model performance is shown in Figure S2). Ultimately, 3,625 genes with probability values greater than 0.7 were sorted out.

Besides the general information such as gene/protein ID, taxonomy ID, general descriptions and orthologous (Figure 2a), FertilityOnline provides high-quality functional annotation information for the collected functional spermatogenic genes. We have classified genes based on developmental stages in spermatogenesis and cell type in testis. Consequently, most of the reported genes were found during meiotic and postmeiotic stages (Table S6), corresponding to spermatocyte and spermatid respectively (Table S7). Additionally, figures collected from references that support functional classification are also displayed on the web. Moreover, we have also provided a manual annotation of gene functions and signaling pathways, and their associated protein complexes are also annotated (Figure 2b).

**Figure 2.**
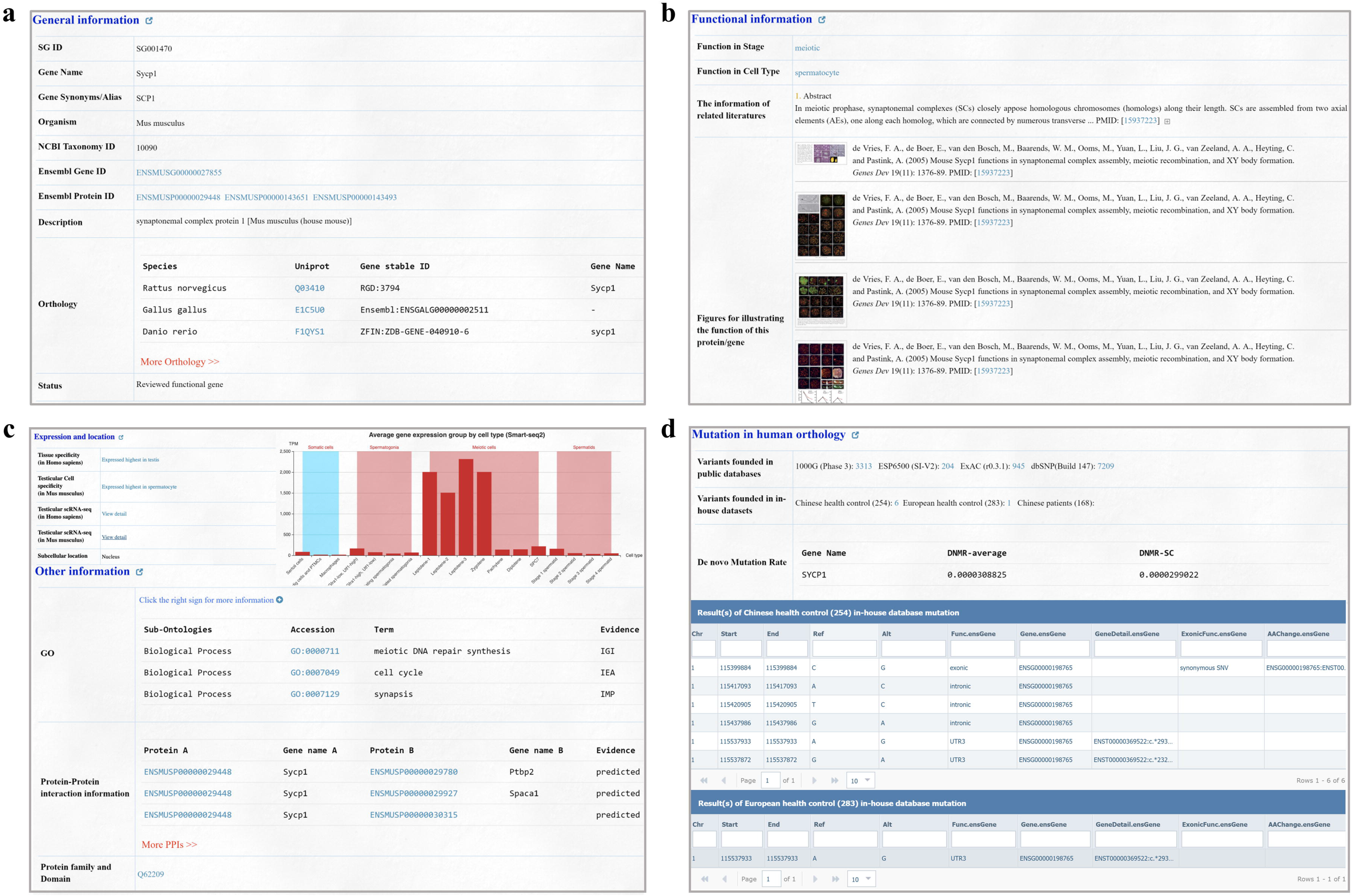
Information integrated in FertilityOnline. (a) A screenshot showing the general information such as gene/protein ID, NCBI taxonomy ID, general descriptions and orthology of the gene (*Sycp1*) used in case study. (b) A screenshot showing the functional information of the gene (*Sycp1*), including functional stage and cells, related literature and figures. (c) The gene expression and location information of the example gene. (d) A screenshot demonstrating the available mutation information of the gene (*Sycp1*) in human orthology. In particular, variants counts in different public databases and our in-house data are provided. Additionally, the statistics of the *de novo* mutation rate of the spermatogenic genes is also shown.

For candidate genes, the references that implicate their function in spermatogenesis, such as information about the reported function, gene expression, protein localization, structure and protein interactions, are included in FertilityOnline. This information will allow users to select candidate genes for experimental validation (Figure 2c).

This database also integrates a range of genetic databases to facilitate screening of pathogenic mutations related to spermatogenesis disorder. In FertilityOnline, users can acquire the counts of variants among different databases and can view the detailed variants information.

*De novo* mutation rate is an important parameter for assessing the pathogenicity of a gene [14, 15]. Generally, genes with higher *de novo* mutation rate appear to be less pathogenic. Therefore, we provided the statistics regarding the *de novo* mutation rate of the spermatogenic genes in FertilityOnline. Users can access this information in mutation section of each page (Figure 2d).

### FertilityOnline Facilitates the Discovery of Functional Genes in Spermatogenesis

Our database provides a feature-rich visual interface for users to screen the genes related to spermatogenesis. Here are some of the functional modules of the web page:

1. ***Search***. Users can search for a specific term, such as the gene/protein name, species, protein complexes, signaling pathways, functional classification and disease characteristics, to find out the gene of interest (Figure S3a).
2. ***Advanced Search***. Users can refine their search results by combining multiple search terms (Figure S3b).
3. ***Browse***. Users can browse all genes that are associated with a certain functional stage, cell type or disease (Figure S3c).
4. ***Blast Search***. By uploading a protein sequence in FASTA format, identical or homologous proteins present in FertilityOnline can be mapped (Figure S3d).
5. ***Homologous Search***. Users can input a gene name and species to obtain the homologous genes in other species. Moreover, they can also select two species and get all the homologous genes (Figure S3e). From the search results, every gene can be further functionally annotated in FertilityOnline (Figure S3f).

### FertilityOnline Facilitates the Discovery of Disease-Causing Variants of Genes Associated within Male Infertility

The major aim of FertilityOnline is to provide a powerful tool to facilitate the screening of disease-causing mutations associated with spermatogenic failure. In FertilityOnline, an analysis module has been provided for users to analyze genes or mutations. After uploading gene or mutation list, the analysis module will annotate it with all available information in FertilityOnline (Figure S4a). To be noted, the uploaded data is temporarily stored on the server and will be automatically deleted after 30 days. The progress of the analysis will be displayed in real-time and on average completes in 5 minutes for a standard VCF format sample (100,000-400,000 variants) (Figure S4b). Finally, the annotation results will be displayed on the web page, and users can filter these results as per their need to sort out candidate genes or mutations (Figure S4c). Moreover, users can perform enrichment analysis using selected genes (Figure S4d) as well as go for further in-depth analysis (Figure S4e). The enriched items are divided into six categories, including functional category, general annotation, gene ontology, disease, pathway and protein domain. A step-by-step protocol described in File S1 and Figure S5.

### Novel *SYCE1* and *STAG3* Mutations Identified by FertilityOnline

Herein, we provide two case studies demonstrating how users can use FertilityOnline to screen the potential pathogenic mutations through the web page. First we uploaded the *.vcf file derived from total exome sequencing data of a Chinese azoospermic patient via the analysis page. FertilityOnline automatically started the entire analysis process and display the progress in real time. Once finished, the complete annotation results for genes and variants were displayed on the web page. Considering the fact that the patient only displayed azoospermia without any other abnormality, the causative gene(s) of this disease were likely to be associated with spermatogenesis. Thus, we set the following parameters in the filter box on the web page: 1) the mutation falls in the exons; 2) the MAF in the 1000G, ESP and ExAc databases is less than 0.05; 3) it is not present in China and Europe with fertility history; 4) the expression level in testis is more than twice than in other tissues; 5) the selection of the reviewed functional genes. With those parameters, a total of 4 mutations in 4 different genes were obtained.

Among them, *Syce1* gene has been reported to be crucial for mouse meiosis, which is consistent with the meiotic arrest phenotype observed in this azoospermia patient (Figure 4a). Thus, the mutations in *SYCE1* are likely the factors causing this patient’s SDA phenotype. The *SYCE1* mutation was further validated by Sanger sequencing (Figure 4b). This caused a nonsense mutation, in which a premature stop codon was introduced at amino acid residue 52 (p.R52*) (Figure 4c), leading to a possible production of a truncated SYCE1 protein in testis. SYCE1 has previously been shown to display aggregates when ectopically expressed in cultured mammalian cells [16]. We took advantage of this observation and examined whether the nonsense mutation of *SYCE1* influences the localization following transfection into Vero cells. Remarkably, WT *SYCE1* expressed aggregates into multiple foci in transfected cells, whereas no foci were observed for mutant *SYCE1* (Figure 4d). Thus, our results suggest that the nonsense mutation of *SYCE1* abrogated the function of *SYCE1*, which is responsible for spermatocyte development arrest in the patient.

**Figure 3.**
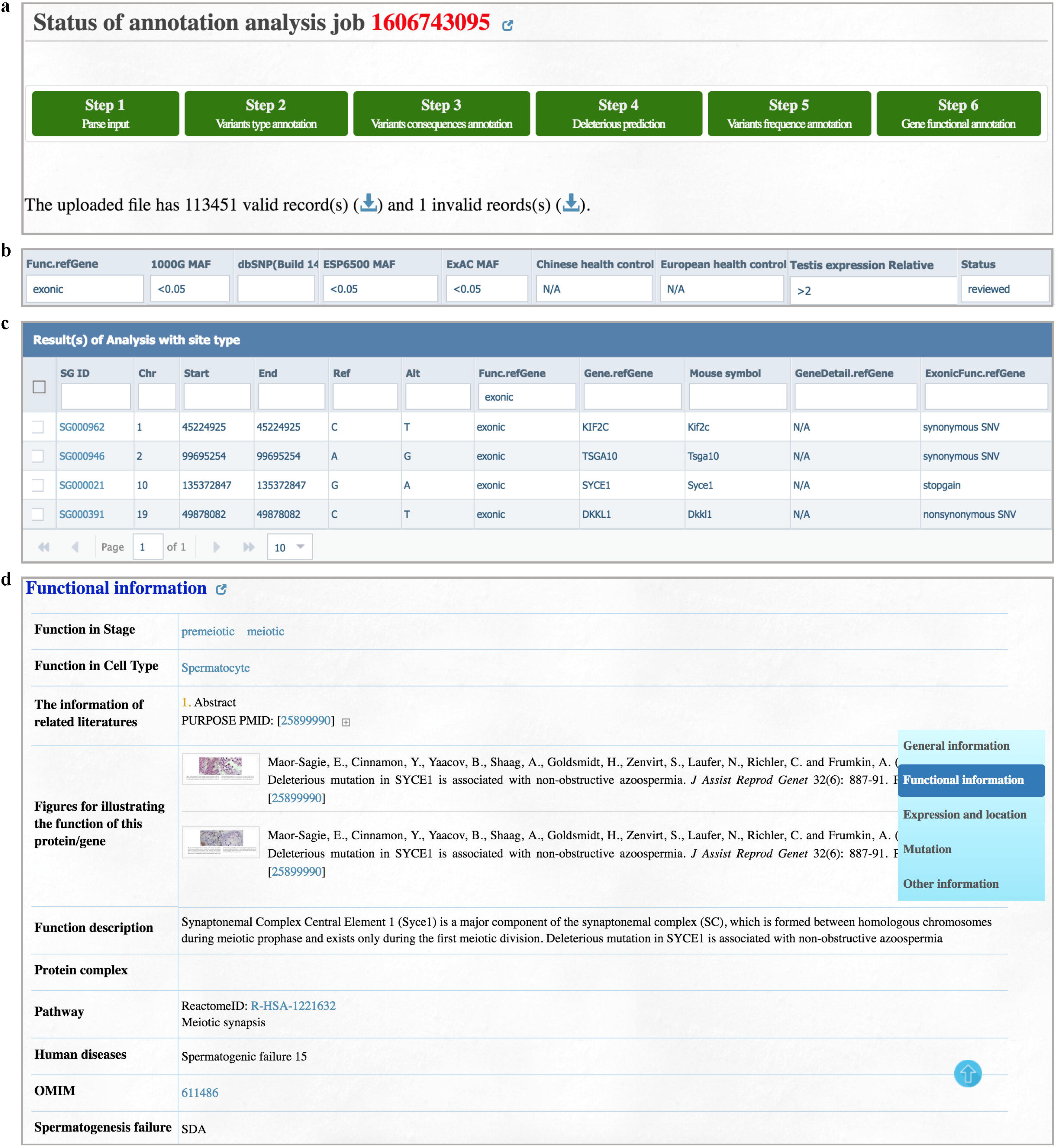
Case studies of how FertilityOnline facilitates the discovery of gene variants. (a) Analysis results of the uploaded *.vcf file containing data from a Chinese azoospermia patient. (b) A representation of the applied filter parameters in the filter box on the web page. (c) Filteration results displaying 4 mutations, corresponding to 4 different genes. (D) Functional information of the candidate gene, *SYCE1*.

**Figure 4.**
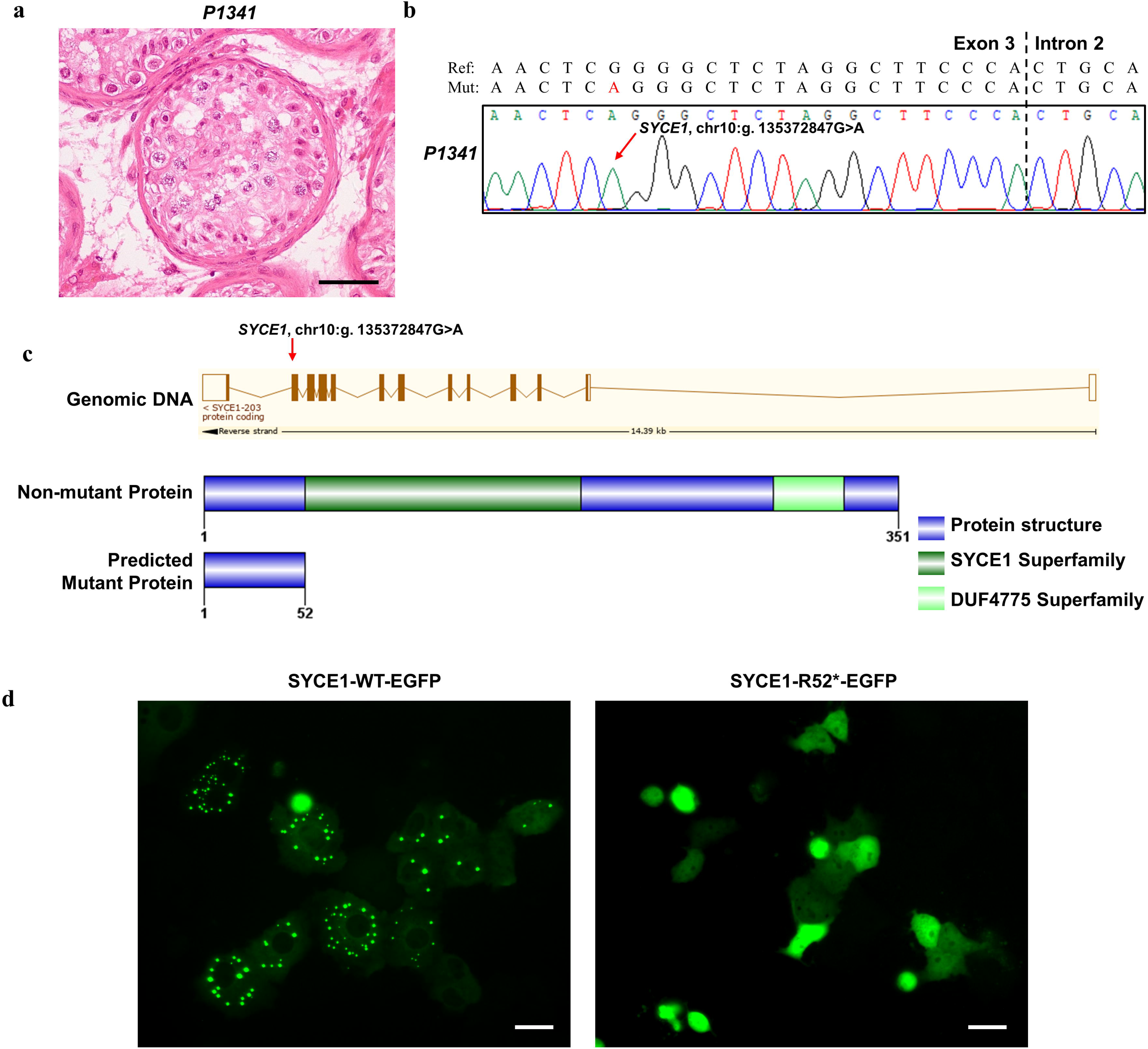
A novel nonsense mutation (c.1634G>A, R52*) in the *SYCE1* gene identified in a male sterile patient by FertilityOnline. (a) Representative images of testicular histology from patient displaying SDA. Scale bar, 50 um. (b) Chromatogram showing the Sanger sequencing result confirming the (*SYCE1*, g. 135372847G>A) mutation in gDNA. A red arrow highlights the mutation site. (c) The exonic map of *SYCE1* is shown in the upper part of the RefSeq transcript (ENST00000368517) showing the position of the novel identified mutations. Verticle boxes are depicted as exons and the line connecting these boxes are introns. The filled boxes represent the coding exone and the non-filled, empty boxes represent the non-coding exons.The non-mutant protein sequence is represented in lower part having 351 AA with 2 domains. The predicted mutant protein length is 52 AA because of the nonsense mutation in the exon 3. (d) Expression pattern of SYCE1 mutational effect. WT SYCE1 has a punctate localization when overexpressed in Vero cells, while, mutant SYCE1 have diffuse localization when overexpressed in Vero cells. Protein expression was analzyed by immunofluorescence microscopy after 36 hours of transfection in Vero cells. Scale bars: 10 μm.

As another example, we uploaded the exome sequencing data from a second Chinese azoospermic patient (Figue 5a and Figure S6a). After getting the annotation results for variants and their carrier genes (Figure S6b), we set parameters in the filter box on the web page (Figure S6c). The pipeline identified 3 mutations in 3 different genes at the end of analysis (Figure S6d). Based on gene information, we focused our attention on STAG3, a component of meiosis specific cohesion complex that is important for meiosis. The *STAG3* mutation was further verified by Sanger sequencing (Figure 5b-c) at both DNA and mRNA levels. Likewise, this mutation also introduced a premature stop codon at residue 357 (p.R357*) (Figure 5d) that possibly produced a C-terminally truncated protein. To confirm this, we generated EGFP-tagged WT *STAG3* and mutant *STAG3* that carried the c.1069C>T in the coding DNA sequence (CDS) and performed the Western blot on cell lysatse following transfection. As expected, the mutant *STAG3* indeed produced a truncated protein at 36kD while the WT STAG3 showed a full-length protein at 134kD (Figure 5e). This evidence validated that c.1069C>T mutation truncated the full-length STAG3 protein at the c-terminal, giving rise to the meiotic arrest in the patient.

**Figure 5.**
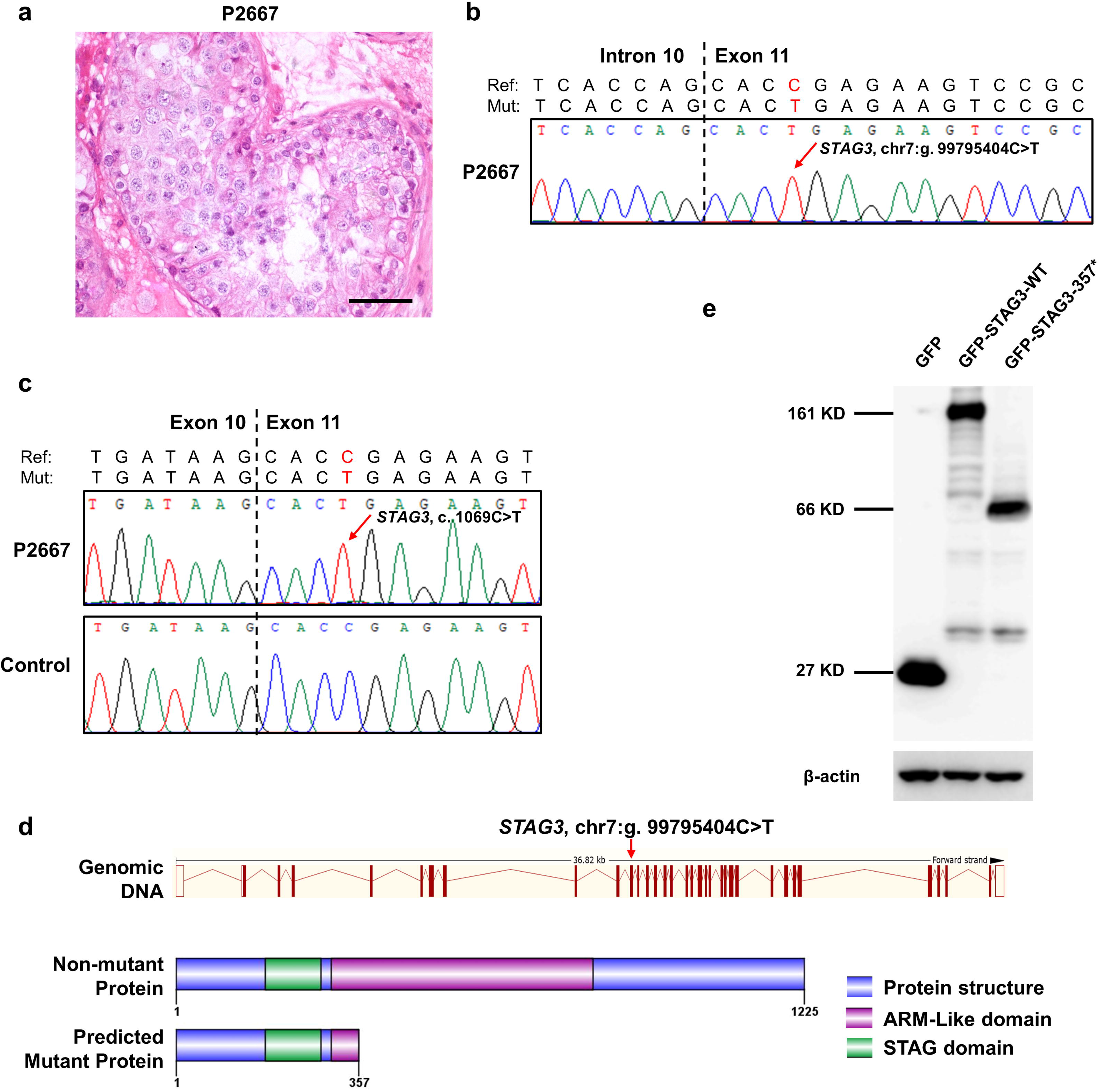
A nonsense mutation (c.1069C>T, R357*) in the *STAG3* gene identified in a male sterile patient by FertilityOnline. (a)Representative sections of testicular histology of patient displaying SDA. Scale bar, 50 um. (b) Chromatogram showing the Sanger resequencing result confirming the (STAG3, g. 99795404C>T) mutation in gDNA. A red arrow highlights the mutation site. (c) Confirmation of *STAG3* (c.1069C>T) mutation at mRNA level. (d) Schematic representation of exons and protein sequence of STAG3. The exonic map of STAG3 is shown in the upper part of the RefSeq transcript (ENST00000426455) showing the position of the novel identified mutations. The WT protein sequence is represented in lower part having 1125 amino acids with 2 domains. The predicted mutant protein length is 357 AA, induced by the nonsense mutation in the exon 11. (e) Western blot shows the protein lysate extracted from cell lines harboring *STAG3* (c.1069C>T) mutation represent ~66□kDa fusion protein corresponding to the predicted truncated STAG3 (~39□kDa) fused to EGFP (~27□kDa). β-actin was used as internal control.

## Discussion

A large number of genes are implicated in the pathogenesis of human diseases, yet the genetic etiology underlying various diseases, e.g., male infertility, remains largely underdetermined [17, 18]. The databases currently available lack depth and accuracy, which makes it difficult to obtain sufficient information to annotate the genes and their mutations. For example, more than two thousand genes that function across different developmental stages of spermatogenesis and in various testicular cell types are involved in production of sperm [6, 7]. Perturbations at any substage during spermatogenesis may eventually lead to infertility, thus the underlying causes of infertility are diverse. Without detailed analysis of the specific phenotype of the abnormality, it is difficult to pinpoint the accurate causative gene and its mutation. The conventional gene annotation databases focus on providing broad-spectrum annotations, so it is not feasible to precisely classify gene functions based on developmental stages or cell types. Therefore, there are urgent needs for specialized database for functional annotation in the field of reproductive biology. Here, the “Functional information” section provided by our database satisfies the aforementioned requirements. FertilityOnline provides not only the detailed functional classification information, but also additional information about genes and diseases. In particular, the phenotypes of genetically modified mice and their corresponding classification to the patient’s “Spermatogenesis failure”, can be examined. With this information, users could readily find out the candidate variants based on the functional information of their carrier genes.

In recent years, WGS and WES are in extensive use to identify candidate pathogenic mutations in an unbiased manner [19, 20], but the number of mutations obtained by WES and WGS is huge. Therefore, integrated information of the expression, localization and function of those genes that carry mutations will greatly help to screen the candidate pathogenic mutations. In this regard, a number of online tools have been developed for variant annotation like MARRVEL, VEP and ANNOVAR [21–23]. Compared to existing tools, FertilityOnline provides more information in detail. First, it contains gene expression information across a panel of tissues and multiple types of cells in testis. This set of information is particulary tailored for genes related to male infertility. For example, if the infertility of a patient is attributed to the meiotic arrest of spermatocytes, most likely the genes with mutation are preferentially or highly expressed in spermatocytes, which allow us to reduce the number of candidate pathogenic genes and mutations for future validation. Second, we have not only provided the general information of gene orthologs across species, but also collected the functional information of these orthologs published in literature. Given that the functions of protein-coding genes are highly conserved and germ cells undergo similar developmental stages between model animals and human, the information provided in our database will facilitate the screening of genes causing male infertility in humans.

Biologists often face the challenge to cope with high-throughput sequencing data. Our attempt to integrate the availabe databases with functional validations through animal models has provided reproductive biologist a systematic module to quickly annotate a list of batch data on their own. In addition, a queuing mechanism was also adopted to allow for the efficient analysis of uploaded tasks from users to ensure timely and stable annotation. For the analyzed results, a screening module is also provided to allow users to reset parameters in the web interface directly, in order to focus on highly likely pathogenic mutations out of a large number of mutations. Furthermore, some links are also provided to help users directly access related databases quickly. For example, during the analyses of the cases presented above, the candidate pathogenic mutations were readily located in *SYCE1* and *STAG3*. To be noted, because we cannot acquire the patients’ testicular tissues to test the existence of mutant mRNAs directly, we cannot rule out the possibility of nonsense-mediated decay for the identified mutations. Instead, we validated the mutations’ effects in cell lines, and found that both mutations affected the protein’s function. Therefore, our database provides an intergrated and systematic platform that allows the batch annotation and screening of gene mutations causing spermatogenic disorders.

## Conclusions

Our database is dedicated to providing a resource for integrating functional gene information regarding spermatogenesis. With this database, users can quickly access the functional information of spermatogenesis-associated genes or dig out candidate disease-causing mutations related to spermatogenic disorders. In particular, this database provides a platform that facilitates the interpretation of the genetic causes of male infertility for diagnosis and research for clinicians as well as biologist.

## Acknowledgments

This project was supported by the National Key Research and Developmental Program of China (2017YFC1001500, 2018YFC1003700, 2016YFC1000600 and 2018YFC1004700), the National Natural Science Foundation of China (31890780, 31630050, 31871514 and 31771668), the Fundamental Research Funds for the Central Universities (YD2070002006).

## Ethical statement

Written informed consent were obtained from the participating subjects and all the human studies are approved by the institutional human ethics committee with the approval number of USTCEC20140003.

## Data availability statement

Data supporting the findings of this study has been deposited in GSA at the National Genomics Data Center under accession number of HRA000257.

## Author’s contribution

J.G, H.Z and A.A constructed the database. D.Z developed the web interface. H.Z and L.J performed the experiments. X.J and A.A wrote the manuscript. Q.B and I.F modified the manuscript. Q.S, Y.Z and X.J conceived and supervised the project.

## Supplementary material

### Supplementary methods

Construction of SVM classifier to predict candidate genes during spermatogenesis

Curation of testicular scRNA-seq data

Performance of variants annotation

### Supplementary figures

Figure S1. Features and the example of predicted results of SVM model

Figure S2. The performance of prediction model

Figure S3. FertilityOnline provides a feature-rich visual interface for users to screen genes related to spermatogenesis

Figure S4. FertilityOnline facilitates the discovery of variants causing male infertility

Figure S5. Step-by-step protocol of variants annotation & filtration

Figure S6. Example of how FertilityOnline facilitates gene and variant analysis

### Supplementary tabless

Table S1. The source of data collected in FertilityOnline

Table S2. Gene expression data collected from ArrayExpress

Table S3. Curated testicular scRNA-seq datasets

Table S4. The performance of variant annotation using FertilityOnline

Table S5. Statistical results of reproted functional genes based on species

Table S6. Statistical results of reported functional genes based on functional stages

Table S7. Statistical results of the reported functional genes based on cell types

### Supplemental figure legends

**Figure S1. Features and the example of predicted results of SVM model.**

Histogram of tissues or cell lines from top 300 features (left) and distribution of the cell types in testes having these 300 features (right); (B) SVM model successfully predicted *Gm4969* as a functional gene in spermatogenesis.

**Figure S2. The performance of prediction model.**

The ROC curve showing the performance of the current dataset (Blue area under the curve (AUC)=0.78).

**Figure S3. FertilityOnline provides a feature-rich visual interface for users to screen genes related to spermatogenesis.**

Users can initiate their search from “Search” option to input query. (b) An advanced search allows users to simultaneously input three terms as query. (c) Browse by species, developmental stage during spermatogenesis, testicular cell types or phenotype in human. (d) BLAST protein sequence search. (e) Browse orthologs for a gene in all species and browse orthologs of all genes in two species. (f) An example of pairwise orthologous browsing in human and mice.

**Figure S4. FertilityOnline facilitates the discovery of variants causing male infertility.**

(a) A user can upload gene or mutation list, the analysis module will annotate it with all available information in FertilityOnline. (b) A displayed progress status of the annotation analysis. (c) A representation of the annotation results on web page that a user can filter to sort out a candidate gene. (d) A web page showing analysis results from a selected batch of genes for the enrichment analysis. (e) Enrichment analysis status with prioritized enriched item based on the functional categories, general annotation, gene ontology, disease, pathway and protein domain.

**Figure S5. Step-by-step protocol of variants annotation & filtration.**

A step-by-step protocol for variants annotation and filtration using a VCF file containing 113,451 variants from the *SYCE1* mutated SDA patient.

(a-b) Prepare the input files. FertilityOnline accepts variants in two formats. (a) VCF format suit for output of regular GATK best practice. The NA columns mean the value of these columns is not necessary. (b) Simple text-based format separated by table which is suitable for several mutation annotation of interest.

(c-e) Quick start. Users can paste the variants into the text form (c) or upload the variants in file (d) with setting the correct reference genome. (e) A processing bar provides a real-time display of analyzing task status.

(f-h) Variants filtration. FertilityOnline provides a rich annotation of genes and mutations, including variant consequence, minor allele frequency (MAF) in 4 public datasets and in-house fertile males, summary of predicted deleterious effects from 13 software and also the rich gene annotation integrated in FertilityOnline. As an example, we made filtration on variants to narrow down the candidates based on the following criteria: (f) MAF < 0.01 in all public datasets (7,778 variants left); (g) Variants located on exonic region (1,339 variants left). (h) Functional gene analysis by setting key words ‘reviewed’ in ‘status’ column and ‘spermatocyte’ in function in cell type column. Finally, the nonsense mutation on *SYCE1* was the most relevant of these 6 mutations, that has the highest probability to cause SDA in the patient.

**Figure S6. Example of how FertilityOnline facilitates gene and variant analysis.**

(a) Analysis results of the uploaded *.vcf file containing data from a Chinese azoospermia patient. (b) A representation of the applied filter parameters in the filter box on the web page. (c) Filtration results displaying 3 mutations in 3 different genes. (d) Functional information of the candidate gene, *STAG3*.

