## Supplementary figures and images for "FertilityOnline, a straight pipeline for functional gene annotation and disease mutation discovery, identifies novel infertility causative mutations in *SYCE1* and *STAG3*"

### Supplementary figure 1

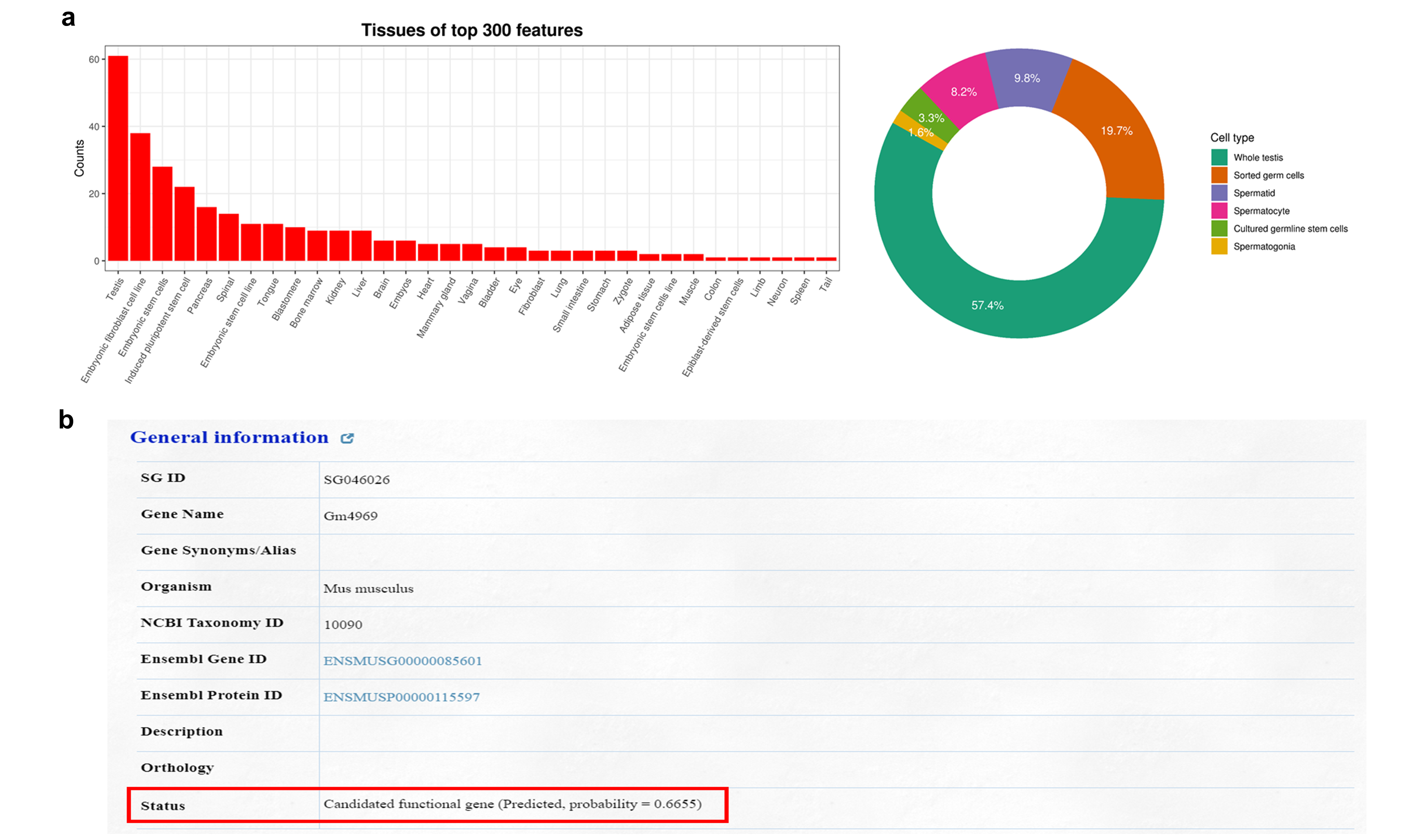

### Supplementary figure 2

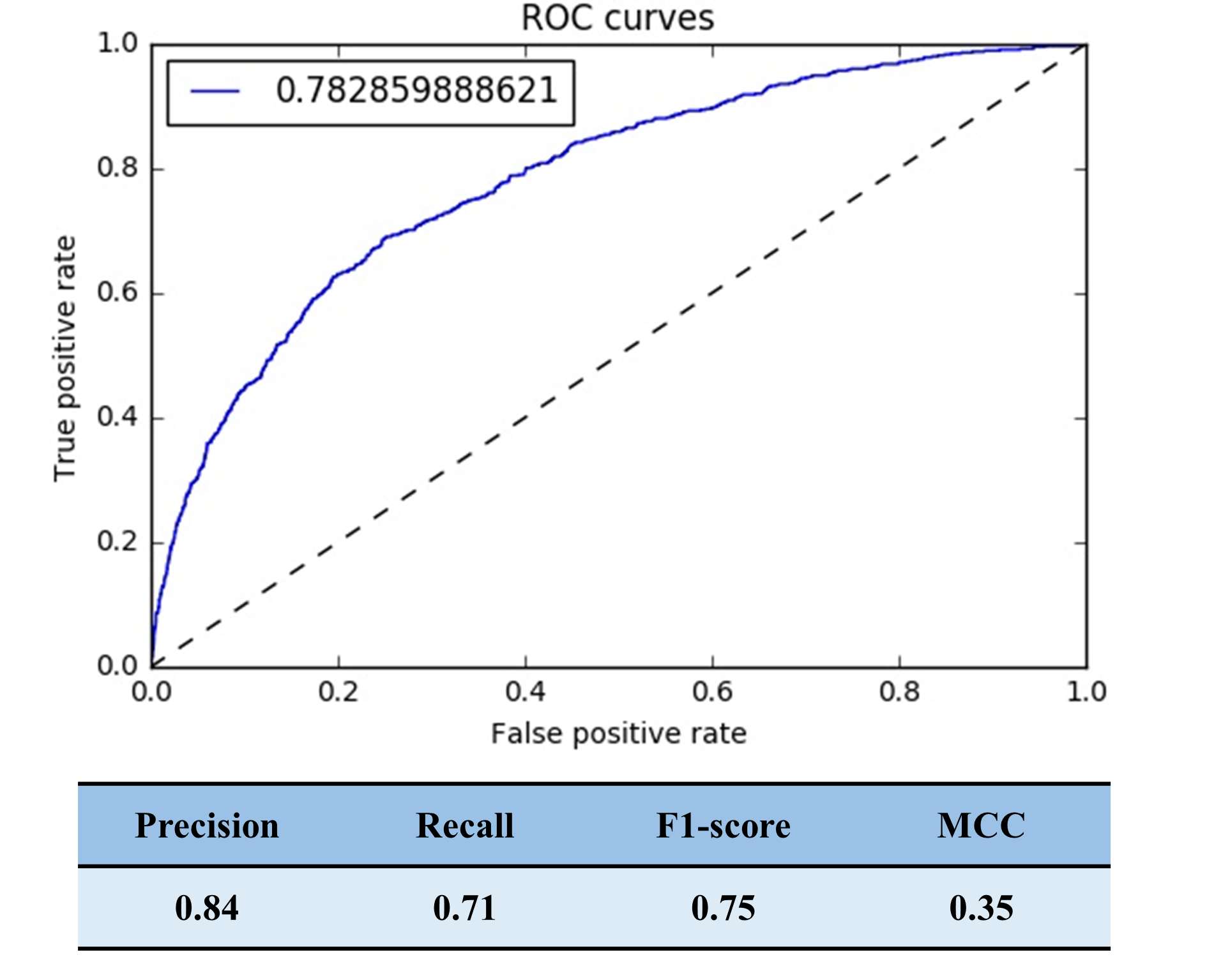

### Supplementary figure 3

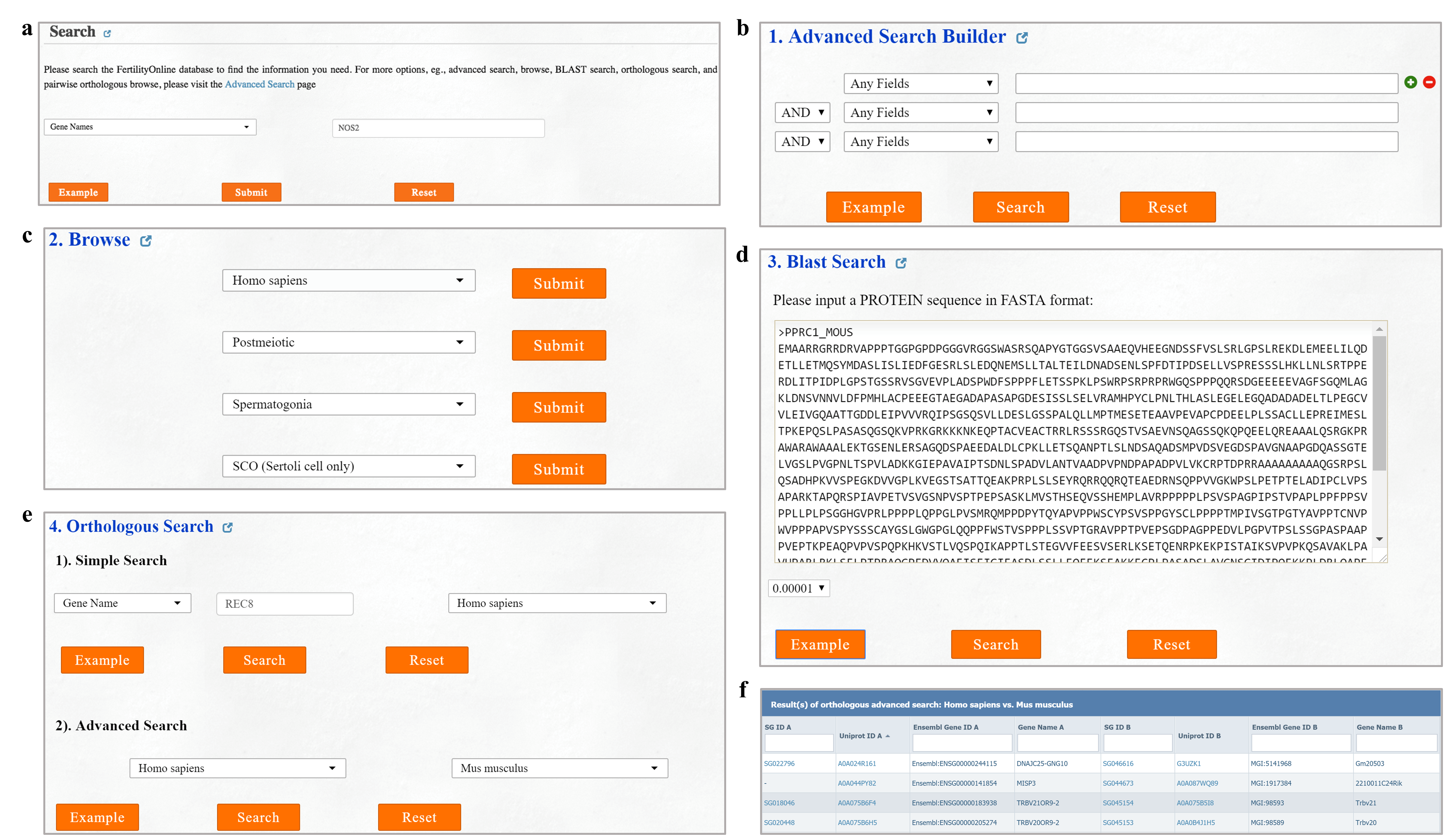

### Supplementary figure 4

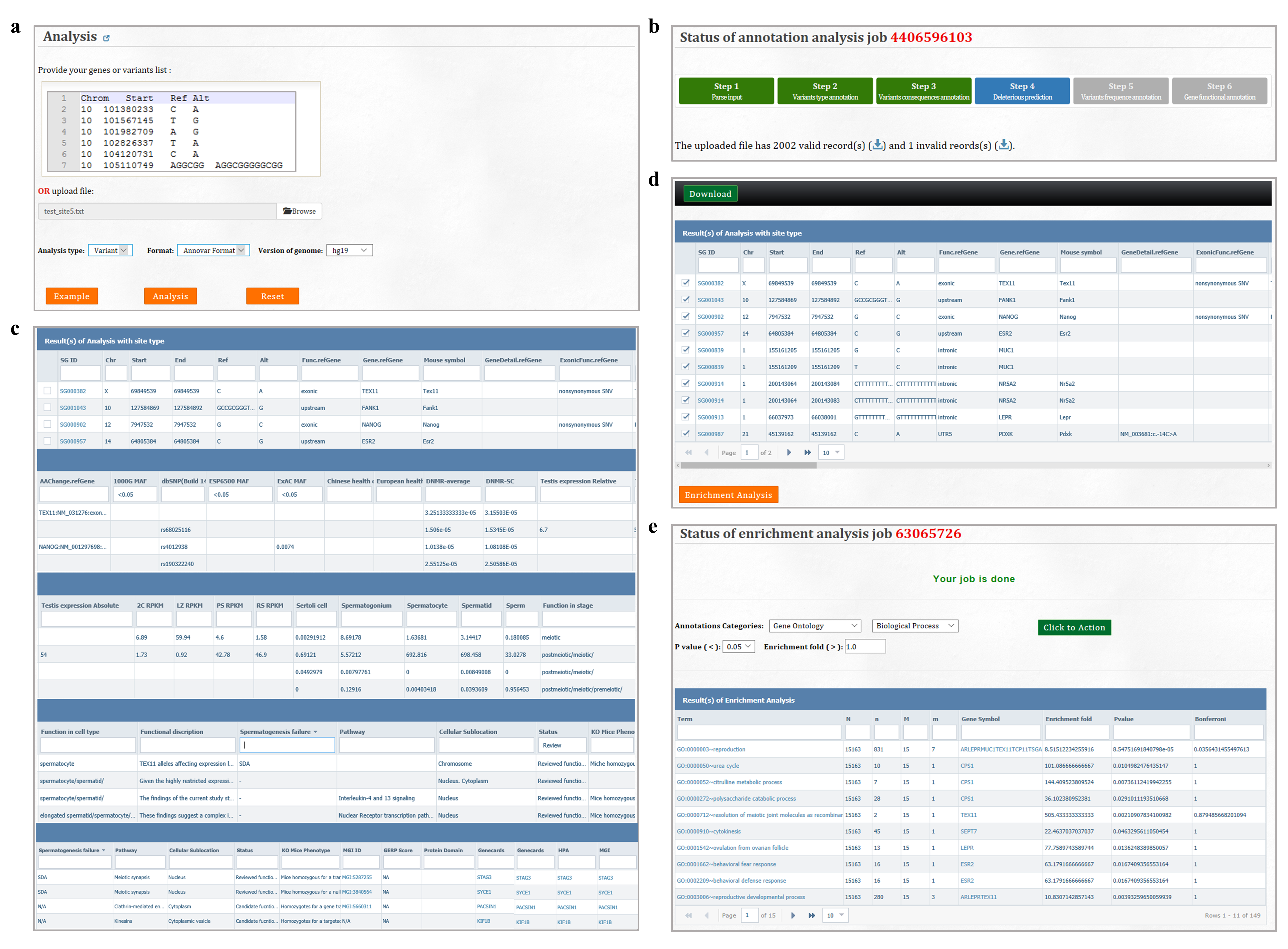

### Supplementary figure 5

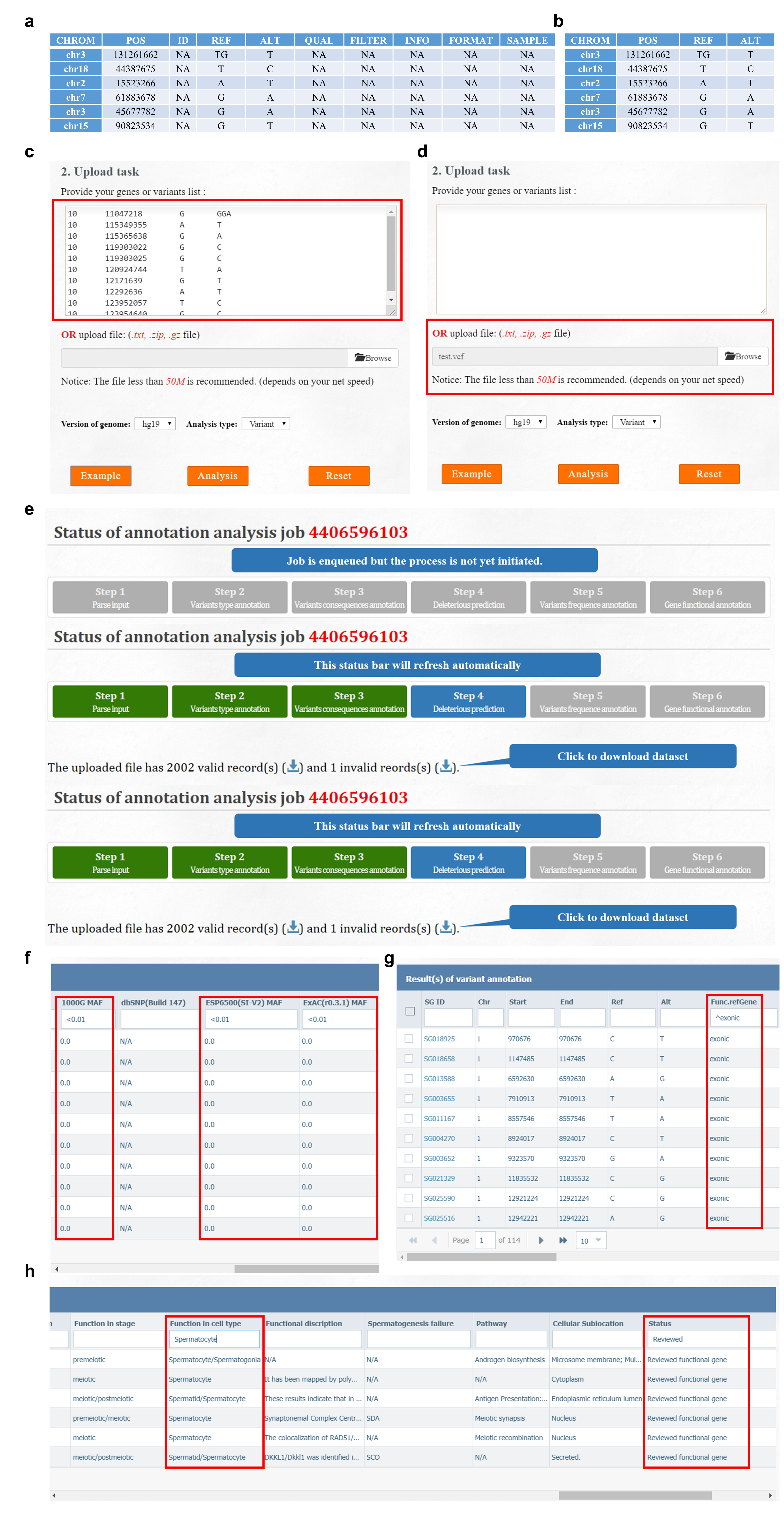

### Supplementary figure 6

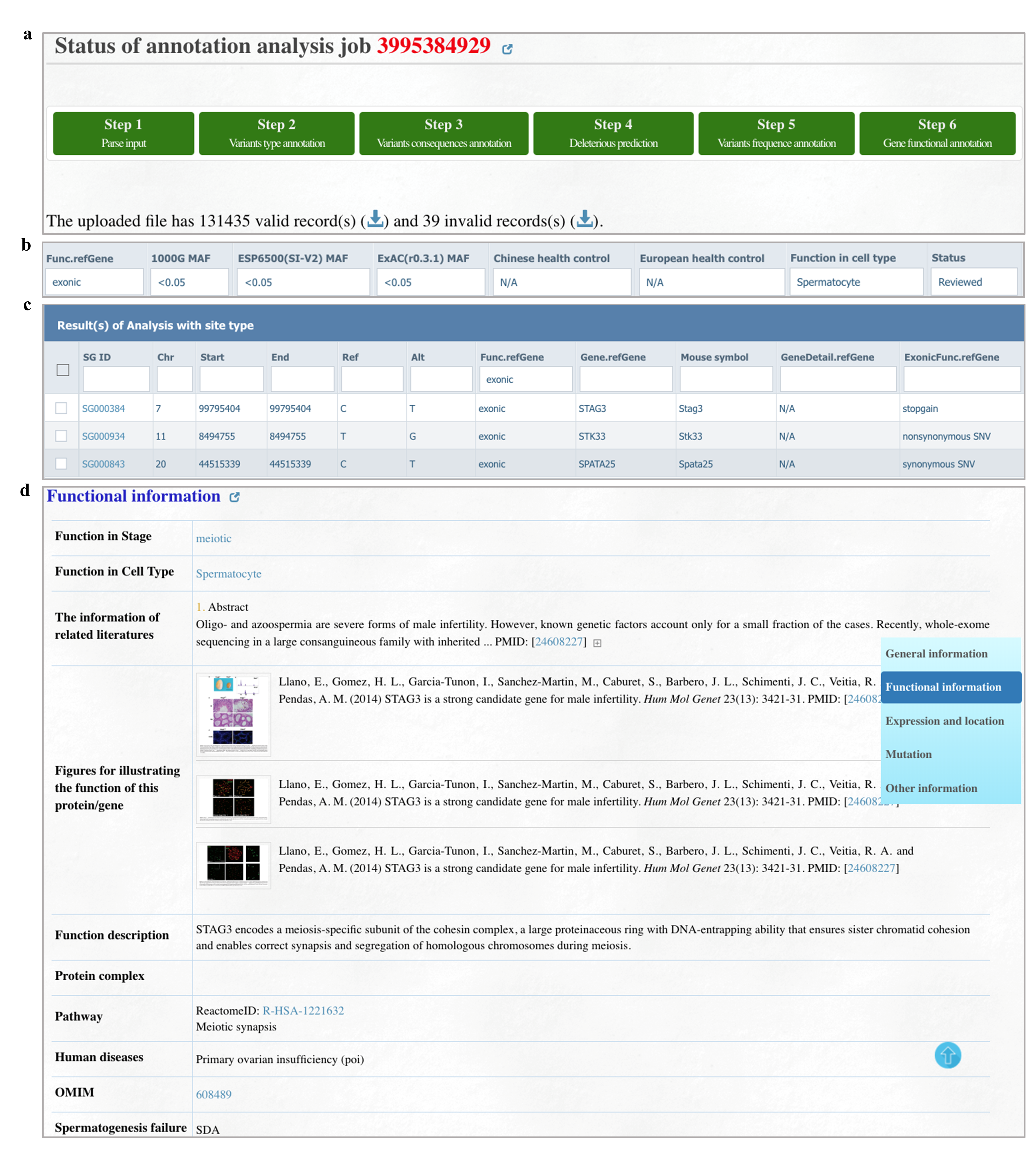
