## Supplementary table 1 for "FertilityOnline, a straight pipeline for functional gene annotation and disease mutation discovery, identifies novel infertility causative mutations in *SYCE1* and *STAG3*"

**Table S1. The source of data collected in FertilityOnline**

| Category | Term | Source |
|---|---|---|
| **General information** | Gene ID | Ensembl (Release 90) |
| | Protein ID | Ensembl (Release 90) |
| | Taxonomy ID | NCBI |
| | Orthology | PANTHER (Release 13.0) |
| **Functional information** | Gene fucntion | Manual curated literatures downloaded from PubMed |
| | Protein complex | CORUM (Released on 02/07/2017) |
| | Protein pathway | DAVID (Version 6.8) |
| | Human diseases | OMIM, ClinVar |
| **Expression and location** | Gene exprssion | ArrayExpress |
| | Subcellular location | NCBI |
| **Mutation** | Public datasets | 1000G (Phase 3) |
| | | ESP6500 (ESP6500SI.V2) |
| | | ExAC (r0.3.1) |
| | | dbSNP (Build 147) |
| | In-house dataset | 254 Chinese health control |
| | | 283 European health control |
| | | 168 Chinese infertile patients |
| | De novo mutation rate | Obtained from work by Jiang *et al.* |
| **Other information** | Gene Ontology | DAVID (Version 6.8) |
| | Protein Domain | DAVID (Version 6.8) |
| | Protein-protein interaction | STRING (Version 10.5) |
