## Supplementary table 2 for "FertilityOnline, a straight pipeline for functional gene annotation and disease mutation discovery, identifies novel infertility causative mutations in *SYCE1* and *STAG3*"

**Table S2. Gene expression data collected from ArrayExpress**

| Accession | Title | Selected assays | Organism | ArrayExpress URL |
|---|---|---|---|---|
| E-GEOD-22004 | Mammalian microRNAs predominantly act to decrease target mRNA levels | 10 | Mus musculus | https://www.ebi.ac.uk/arrayexpress/experiments/E-GEOD-22004/ |
| E-GEOD-36114 | Genome-wide analysis of histone modification, protein-DNA binding, cytosine methylation and transcriptome data in mouse and human ES cells and pig iPS cells | 2 | Mus musculus | https://www.ebi.ac.uk/arrayexpress/experiments/E-GEOD-36114/ |
| E-GEOD-36294 | High-throughput sequencing of sequentially reprogrammed iPS cells reveals key epigenetic modifications correlated with reduced pluripotency of iPS cells | 52 | Mus musculus | https://www.ebi.ac.uk/arrayexpress/experiments/E-GEOD-36294/ |
| E-GEOD-38371 | Latent enhancers unveiled by stimulation expand and adapt the available cis-regulatory repertoire (RNA-seq) | 28 | Mus musculus | https://www.ebi.ac.uk/arrayexpress/experiments/E-GEOD-38371/ |
| E-GEOD-38495 | Full-length mRNA-Seq from single-cell levels of RNA and individual circulating tumor | 29 | Mus musculus | https://www.ebi.ac.uk/arrayexpress/experiments/E-GEOD-38495/ |
| E-GEOD-40463 | STATs Shape the Active Enhancer Landscape of T Cell Populations | 2 | Mus musculus | https://www.ebi.ac.uk/arrayexpress/experiments/E-GEOD-40463/ |
| E-GEOD-41710 | Global gene expression analysis of Dot1l-deficient and control intestinal villus cells in mouse | 1 | Mus musculus | https://www.ebi.ac.uk/arrayexpress/experiments/E-GEOD-41710/ |
| E-GEOD-41785 | Global analysis of Upf1 in mESCs reveals expanded scope of nonsense-mediated mRNA decay | 3 | Mus musculus | https://www.ebi.ac.uk/arrayexpress/experiments/E-GEOD-41785/ |
| E-GEOD-42268 | Quartz-Seq: a simple and highly quantitative method for single-cell RNA-Seq | 60 | Mus musculus | https://www.ebi.ac.uk/arrayexpress/experiments/E-GEOD-42268/ |
| E-GEOD-42999 | ?3 agonist administration before reperfusion reduces infarct size and improves long term cardiac function in small and large animal models of myocardial infarction | 6 | Mus musculus | https://www.ebi.ac.uk/arrayexpress/experiments/E-GEOD-42999/ |
| E-GEOD-43013 | Gene Expression Defines Natural Changes in Mammalian Lifespan | 9 | Mus musculus | https://www.ebi.ac.uk/arrayexpress/experiments/E-GEOD-43013/ |
| E-GEOD-43970 | Reconstruction of the dynamic regulatory network that controls Th17 cell differentiation by systematic perturbation in primary cells | 32 | Mus musculus | https://www.ebi.ac.uk/arrayexpress/experiments/E-GEOD-43970/ |
| E-GEOD-45597 | RNA sequencing of bone marrow hematopoietic progenitor and Gr1+/Mac1+ cells derived from Diap1 wild type, heterozygous, and knockout mice | 6 | Mus musculus | https://www.ebi.ac.uk/arrayexpress/experiments/E-GEOD-45597/ |
| E-GEOD-45719 | Single-cell RNA-Seq reveals dynamic, random monoallelic gene expression in mammalian cells | 292 | Mus musculus | https://www.ebi.ac.uk/arrayexpress/experiments/E-GEOD-45719/ |
| E-GEOD-45982 | EZH2 is required for germinal center formation and somatic EZH2 mutations promote lymphoid transformation | 6 | Mus musculus | https://www.ebi.ac.uk/arrayexpress/experiments/E-GEOD-45982/ |
| E-GEOD-46664 | Digital Gene Expression Tag Profiling after Wt1 deletion in Sertoli cell | 2 | Mus musculus | https://www.ebi.ac.uk/arrayexpress/experiments/E-GEOD-46664/ |
| E-GEOD-46700 | AID stabilizes a stem cell phenotype by removing epigenetic memory of secondary pluripotency network genes | 4 | Mus musculus | https://www.ebi.ac.uk/arrayexpress/experiments/E-GEOD-46700/ |
| E-GEOD-46716 | Indistinguishable Nucleosome Organizations in Embryonic Stem Cells (ESCs) and Induced Pluripotent Stem Cells (iPSCs) Reprogrammed from Somatic Cells Belonging to Three Different Germ Layers[RNA-seq] | 8 | Mus musculus | https://www.ebi.ac.uk/arrayexpress/experiments/E-GEOD-46716/ |
| E-GEOD-47454 | Transcriptome Sequencing (RNA-seq) of Ara-C Resistant Murine AML Cell Lines Identifies Mechanisms of Resistance | 4 | Mus musculus | https://www.ebi.ac.uk/arrayexpress/experiments/E-GEOD-47454/ |
| E-GEOD-47537 | Gata6 | 8 | Mus musculus | https://www.ebi.ac.uk/arrayexpress/experiments/E-GEOD-47537/ |
| E-GEOD-48206 | Gene Expression Profile for knocking down Tet1 in pre-iPSCs culturing in Vc medium | 6 | Mus musculus | https://www.ebi.ac.uk/arrayexpress/experiments/E-GEOD-48206/ |
| E-GEOD-48419 | Gene expression analysis of Ash1l mutant mouse embryonic stem cells | 4 | Mus musculus | https://www.ebi.ac.uk/arrayexpress/experiments/E-GEOD-48419/ |
| E-GEOD-48480 | Oncogenic Hras-dependent epidermal growth is inhibited following RNAi-mediated depletion of Ctnnb1 and Mllt6 | 5 | Mus musculus | https://www.ebi.ac.uk/arrayexpress/experiments/E-GEOD-48480/ |
| E-GEOD-48654 | Genes regulated in mouse liver by expressing ectopically hURI in hepatocytes | 1 | Mus musculus | https://www.ebi.ac.uk/arrayexpress/experiments/E-GEOD-48654/ |
| E-GEOD-48974 | RNA-Seq of eye tissues from C57BL/6J background mice at 1.5 h and 9.0 h after light | 5 | Mus musculus | https://www.ebi.ac.uk/arrayexpress/experiments/E-GEOD-48974/ |
| E-GEOD-49321 | Improved Smart-Seq for sensitive full-length transcriptome profiling in single cells | 76 | Mus musculus | https://www.ebi.ac.uk/arrayexpress/experiments/E-GEOD-49321/ |
| E-GEOD-49624 | Chromatin and Transcription Transitions of Mammalian Adult Germline Stem Cells and Spermatogenesis | 6 | Mus musculus | https://www.ebi.ac.uk/arrayexpress/experiments/E-GEOD-49624/ |
| E-GEOD-49931 | The transcription factor IRF4 is essential for T cell receptor affinity mediated metabolic programming and clonal expansion of T cells | 10 | Mus musculus | https://www.ebi.ac.uk/arrayexpress/experiments/E-GEOD-49931/ |
| E-GEOD-49949 | Loss of Sip1 leads to migration defects and retention of ectodermal markers during lens development | 3 | Mus musculus | https://www.ebi.ac.uk/arrayexpress/experiments/E-GEOD-49949/ |
| E-GEOD-51011 | Selective transcriptional regulation by Myc in cellular growth control and | 11 | Mus musculus | https://www.ebi.ac.uk/arrayexpress/experiments/E-GEOD-51011/ |
| E-GEOD-51281 | EGFRvIII/STAT3 targets in glioblastoma pathogenesis | 4 | Mus musculus | https://www.ebi.ac.uk/arrayexpress/experiments/E-GEOD-51281/ |
| E-GEOD-52069 | Maternal hematopoietic TNF, via milk chemokines, programs hippocampal development and memory | 2 | Mus musculus | https://www.ebi.ac.uk/arrayexpress/experiments/E-GEOD-52069/ |
| E-GEOD-52396 | C/EBP? poises B cells for rapid reprogramming into iPS cells [RNA-Seq] | 2 | Mus musculus | https://www.ebi.ac.uk/arrayexpress/experiments/E-GEOD-52396/ |
| E-GEOD-52583 | High throughput quantitative whole transcriptome analysis of distal mouse lung epithelial cells from various developmental stages (E14.5, E16.5, E18.5 and adult) | 11 | Mus musculus | https://www.ebi.ac.uk/arrayexpress/experiments/E-GEOD-52583/ |
| E-GEOD-52801 | Assessing the ceRNA hypothesis with quantitative measurements of miRNA and target abundance | 19 | Mus musculus | https://www.ebi.ac.uk/arrayexpress/experiments/E-GEOD-52801/ |
| E-GEOD-53150 | RNAseq analysis of mature thymic NKT cells and immature thymic DN1eP cells | 25 | Mus musculus | https://www.ebi.ac.uk/arrayexpress/experiments/E-GEOD-53150/ |
| E-GEOD-53212 | Nanog Independent Reprogramming to iPSCs with Canonical Factors | 16 | Mus musculus | https://www.ebi.ac.uk/arrayexpress/experiments/E-GEOD-53212/ |
| E-GEOD-53297 | Adult stem cells in the small intestine are intrinsically programmed with their location-specific function | 3 | Mus musculus | https://www.ebi.ac.uk/arrayexpress/experiments/E-GEOD-53297/ |
| E-GEOD-54154 | RNA-seq analysis of diabetes induced changes in macrophage transcriptome | 4 | Mus musculus | https://www.ebi.ac.uk/arrayexpress/experiments/E-GEOD-54154/ |
| E-GEOD-54412 | Investigation of the role of histone modification propagating activity of GLP | 5 | Mus musculus | https://www.ebi.ac.uk/arrayexpress/experiments/E-GEOD-54412/ |
| E-GEOD-54461 | Temporal gene expression across osteoblastogenesis | 27 | Mus musculus | https://www.ebi.ac.uk/arrayexpress/experiments/E-GEOD-54461/ |
| E-GEOD-55291 | Noncoding RNA transcriptome analysis during cellular reprogramming | 94 | Mus musculus | https://www.ebi.ac.uk/arrayexpress/experiments/E-GEOD-55291/ |
| E-GEOD-55344 | Temporal dynamics and developmental memory of 3D chromatin architecture at Hox gene loci | 2 | Mus musculus | https://www.ebi.ac.uk/arrayexpress/experiments/E-GEOD-55344/ |
| E-GEOD-55698 | Variant PRC1 complex dependent H2A ubiquitylation drives PRC2 recruitment and polycomb domain formation | 6 | Mus musculus | https://www.ebi.ac.uk/arrayexpress/experiments/E-GEOD-55698/ |
| E-GEOD-55800 | Translational profiling of hypothalamic and midbrain neurons that project to the nucleus accumbens | 6 | Mus musculus | https://www.ebi.ac.uk/arrayexpress/experiments/E-GEOD-55800/ |
| E-GEOD-55870 | Gene Expression Analysis of Cancer-Associated Fibroblast (CAF) compared to Normal Fibroblast (NF) [RNA-seq] | 2 | Mus musculus | https://www.ebi.ac.uk/arrayexpress/experiments/E-GEOD-55870/ |
| E-GEOD-56138 | Reorganization of enhancer patterns in transition from naïve to primed pluripotency | 9 | Mus musculus | https://www.ebi.ac.uk/arrayexpress/experiments/E-GEOD-56138/ |
| E-GEOD-56575 | Rapid proliferation and differentiation impairs the development of memory CD8+ T cells in early life | 3 | Mus musculus | https://www.ebi.ac.uk/arrayexpress/experiments/E-GEOD-56575/ |
| E-GEOD-56697 | Programming and inheritance of parental DNA methylomes in mammals | 1 | Mus musculus | https://www.ebi.ac.uk/arrayexpress/experiments/E-GEOD-56697/ |
| E-GEOD-56762 | Histone variant H3.3 is an essential maternal factor for oocyte reprogramming | 5 | Mus musculus | https://www.ebi.ac.uk/arrayexpress/experiments/E-GEOD-56762/ |
| E-GEOD-56932 | Epigenetic and transcriptional regulation of starvation-induced atrophy and autophagy programs by Foxk1 and Sin3A | 2 | Mus musculus | https://www.ebi.ac.uk/arrayexpress/experiments/E-GEOD-56932/ |
| E-GEOD-56986 | Tet proteins in DNA demethylation and cell fate restriction | 2 | Mus musculus | https://www.ebi.ac.uk/arrayexpress/experiments/E-GEOD-56986/ |
| E-GEOD-57313 | MicroRNAs Shape Circadian Hepatic Gene Expression on a Transcriptome-Wide Scale | 13 | Mus musculus | https://www.ebi.ac.uk/arrayexpress/experiments/E-GEOD-57313/ |
| E-GEOD-57577 | Engineering of a histone-recognition domain in Dnmt3a alters the epigenetic landscape of ESCs revealing changes in lineage specification and chromosomal stability | 8 | Mus musculus | https://www.ebi.ac.uk/arrayexpress/experiments/E-GEOD-57577/ |
| E-GEOD-57959 | Comparison of expression data of 2 & 4- months old CRYAAN101D versus CRYAAWT lenses | 4 | Mus musculus | https://www.ebi.ac.uk/arrayexpress/experiments/E-GEOD-57959/ |
| E-GEOD-58006 | Changes in nucleosome occupancy associated with metabolic alterations in aged mammalian liver | 3 | Mus musculus | https://www.ebi.ac.uk/arrayexpress/experiments/E-GEOD-58006/ |
| E-GEOD-58398 | The histone lysine demethylase Kdm6b is required for activity-dependent preconditioning of hippocampal neuronal survival | 4 | Mus musculus | https://www.ebi.ac.uk/arrayexpress/experiments/E-GEOD-58398/ |
| E-GEOD-58429 | RNA-seq studies reveal new insights into p63 and the transcriptomic landscape of the mouse skin | 3 | Mus musculus | https://www.ebi.ac.uk/arrayexpress/experiments/E-GEOD-58429/ |
| E-GEOD-58669 | A Dach2-Hdac9-Myog-Gdf5 signaling system regulates regeneration of neuromuscular synapses | 1 | Mus musculus | https://www.ebi.ac.uk/arrayexpress/experiments/E-GEOD-58669/ |
| E-GEOD-58833 | Effect of Bicaudal C1 deletion on E13.5 mouse pancreatic mRNA profile | 6 | Mus musculus | https://www.ebi.ac.uk/arrayexpress/experiments/E-GEOD-58833/ |
| E-GEOD-58998 | Transcription profiling by high throughput sequencing of mice with a partial or complete deletion of Ezh2 in regulatory T cells | 11 | Mus musculus | https://www.ebi.ac.uk/arrayexpress/experiments/E-GEOD-58998/ |
| E-GEOD-59119 | The contribution of cohesin-SA1 to chromatin architecture and gene expression in two murine tissues [RNA-seq] | 2 | Mus musculus | https://www.ebi.ac.uk/arrayexpress/experiments/E-GEOD-59119/ |
| E-GEOD-59215 | Transcriptome impact of acute deletion of Gata3 in murine pro-T cells | 2 | Mus musculus | https://www.ebi.ac.uk/arrayexpress/experiments/E-GEOD-59215/ |
| E-GEOD-59285 | The G Protein-coupled Receptor P2Y14 Influences Insulin Release and Smooth Muscle Function in Mice | 20 | Mus musculus | https://www.ebi.ac.uk/arrayexpress/experiments/E-GEOD-59285/ |
| E-GEOD-59777 | Escape from X inactivation in mouse tissues (RNA-Seq) | 1 | Mus musculus | https://www.ebi.ac.uk/arrayexpress/experiments/E-GEOD-59777/ |
| E-GEOD-59833 | Transcriptional Pause Release Is a Rate-Limiting Step for Somatic Cell Reprogramming | 1 | Mus musculus | https://www.ebi.ac.uk/arrayexpress/experiments/E-GEOD-59833/ |
| E-GEOD-60231 | RNA-Seq analysis to profile the transcriptome of Bone Marrow Dendritic Cells exposed to different antigen delivery systems | 8 | Mus musculus | https://www.ebi.ac.uk/arrayexpress/experiments/E-GEOD-60231/ |
| E-GEOD-60969 | Gcn5 and PCAF negatively regulate interferon ? production through HAT-independent inhibition of TBK1 | 5 | Mus musculus | https://www.ebi.ac.uk/arrayexpress/experiments/E-GEOD-60969/ |
| E-GEOD-61095 | OGG1-initiated DNA base excision repair is linked to inflammatory gene expression and lung inflammation | 1 | Mus musculus | https://www.ebi.ac.uk/arrayexpress/experiments/E-GEOD-61095/ |
| E-GEOD-61300 | High throughput quantitative whole transcriptome analysis of CC10- B4+ and Krt5-CreERT2 - labeled distal lung cells | 1 | Mus musculus | https://www.ebi.ac.uk/arrayexpress/experiments/E-GEOD-61300/ |
| E-GEOD-61331 | Role of Tet3 and DNA replication in zygotic demethylation of both paternal and maternal genomes | 8 | Mus musculus | https://www.ebi.ac.uk/arrayexpress/experiments/E-GEOD-61331/ |
| E-GEOD-61486 | Lineage Reprogramming of mouse fibroblasts into induced cardiac progenitor cells by defined factors | 7 | Mus musculus | https://www.ebi.ac.uk/arrayexpress/experiments/E-GEOD-61486/ |
| E-GEOD-61636 | Transcription profiling by high throughput sequencing of murine bone marrow endothelial cells and bone marrow stroma, in vitro and in vivo, with and without hematopoietic stem cells co-culture | 24 | Mus musculus | https://www.ebi.ac.uk/arrayexpress/experiments/E-GEOD-61636/ |
| E-GEOD-61694 | MEF OSK O KS T20 | 5 | Mus musculus | https://www.ebi.ac.uk/arrayexpress/experiments/E-GEOD-61694/ |
| E-GEOD-61887 | Activity-Induced DNA Breaks Govern the Expression of Neuronal Early-Response | 2 | Mus musculus | https://www.ebi.ac.uk/arrayexpress/experiments/E-GEOD-61887/ |
| E-GEOD-62310 | RNA-SEQ analysis of the developing lower urogenital region | 10 | Mus musculus | https://www.ebi.ac.uk/arrayexpress/experiments/E-GEOD-62310/ |
| E-GEOD-62432 | Comparative transcriptomic analysis of self-organized, in vitro generated optic tissues | 15 | Mus musculus | https://www.ebi.ac.uk/arrayexpress/experiments/E-GEOD-62432/ |
| E-GEOD-62633 | Transcription Factors GATA4 and HNF4A Control Distinct Aspects of Intestinal Homeostasis in Conjunction With the Transcription Factor CDX2 | 4 | Mus musculus | https://www.ebi.ac.uk/arrayexpress/experiments/E-GEOD-62633/ |
| E-GEOD-63137 | Epigenomic Signatures of Neuronal Diversity in the Mammalian Brain | 8 | Mus musculus | https://www.ebi.ac.uk/arrayexpress/experiments/E-GEOD-63137/ |
| E-GEOD-63751 | Next Generation Sequencing Facilitates Quantitative Analysis of Bone Marrow Macrophages and Spleenic Macrophages Transcriptomes in mouse T cell acute lymphoblastic leukemia | 2 | Mus musculus | https://www.ebi.ac.uk/arrayexpress/experiments/E-GEOD-63751/ |
| E-GEOD-64040 | Functional and Mechanistic Studies of XPC DNA-Repair Complex as Transcriptional Coactivator in Embryonic Stem Cells | 3 | Mus musculus | https://www.ebi.ac.uk/arrayexpress/experiments/E-GEOD-64040/ |
| E-GEOD-65031 | 8-Oxoguanine DNA glycosylase-1 DNA repair-signaling induces gene expression associated to airway remodeling | 3 | Mus musculus | https://www.ebi.ac.uk/arrayexpress/experiments/E-GEOD-65031/ |
| E-GEOD-65337 | RNA-sequencing of Postnatal Day 10 Wild-type and Nfix KO Subventricular Zone-derived Primary Monolayer-cultured Neural Stem Cells | 3 | Mus musculus | https://www.ebi.ac.uk/arrayexpress/experiments/E-GEOD-65337/ |
| E-GEOD-65344 | PU.1 regulates T-lineage gene expression and progression via indirect repression during early T-cell development | 4 | Mus musculus | https://www.ebi.ac.uk/arrayexpress/experiments/E-GEOD-65344/ |
| E-GEOD-65636 | Cellular response to heat shock and ER stress | 1 | Mus musculus | https://www.ebi.ac.uk/arrayexpress/experiments/E-GEOD-65636/ |
| E-GEOD-65665 | Gene expression profiling of effect of Yap inhibition in a genetically engineered mouse model of hepatocellular carcinoma | 7 | Mus musculus | https://www.ebi.ac.uk/arrayexpress/experiments/E-GEOD-65665/ |
| E-GEOD-66092 | PAPERCLIP identifies novel microRNA targets and provides insights into mRNA alternative polyadenylation | 3 | Mus musculus | https://www.ebi.ac.uk/arrayexpress/experiments/E-GEOD-66092/ |
| E-GEOD-66147 | The PIAS-like coactivator Zmiz1 directly and selectively coregulates Notch1 in T-cell development and leukemia | 12 | Mus musculus | https://www.ebi.ac.uk/arrayexpress/experiments/E-GEOD-66147/ |
| E-GEOD-66202 | Intrinsic age-dependent changes and cell-cell contacts regulate nephron progenitor | 23 | Mus musculus | https://www.ebi.ac.uk/arrayexpress/experiments/E-GEOD-66202/ |
| E-GEOD-66231 | Effect of pharmacological inhibition of ASBT on bile composition and sclerosing cholangitis in mdr2 knockout mice | 2 | Mus musculus | https://www.ebi.ac.uk/arrayexpress/experiments/E-GEOD-66231/ |
| E-GEOD-66440 | IGF2BP2/IMP2 Deficient Mice Resist Obesity through enhanced translation of Ucp1 mRNA and other mRNAs encoding Mitochondrial Proteins | 14 | Mus musculus | https://www.ebi.ac.uk/arrayexpress/experiments/E-GEOD-66440/ |
| E-GEOD-66557 | Transcriptome analyses of skeletal muscle in ?B-crystallin/HspB2 knockout and wild-type mice on a normal or high fat diet | 12 | Mus musculus | https://www.ebi.ac.uk/arrayexpress/experiments/E-GEOD-66557/ |
| E-GEOD-66736 | Tex10 Coordinates Epigenetic Control of Super-Enhancer Activity for Pluripotency and Reprogramming | 2 | Mus musculus | https://www.ebi.ac.uk/arrayexpress/experiments/E-GEOD-66736/ |
| E-GEOD-66744 | Decoding breast cancer tissue-stroma interactions using species-specific sequencing | 21 | Mus musculus | https://www.ebi.ac.uk/arrayexpress/experiments/E-GEOD-66744/ |
| E-GEOD-67009 | ESRP2 Regulates A Conserved And Cell-Type-Specific Splicing Program to Support Postnatal Liver Maturation | 6 | Mus musculus | https://www.ebi.ac.uk/arrayexpress/experiments/E-GEOD-67009/ |
| E-GEOD-67052 | RNA-sequencing of DYRK1A-deficient (CKO) pre-B and pre-T cells | 2 | Mus musculus | https://www.ebi.ac.uk/arrayexpress/experiments/E-GEOD-67052/ |
| E-GEOD-67123 | Tracing the formation of hematopoietic stem cells in mouse embryos by single-cell functional and RNA-Seq analyses | 66 | Mus musculus | https://www.ebi.ac.uk/arrayexpress/experiments/E-GEOD-67123/ |
| E-GEOD-67207 | Effect of estrogen and selective estrogen receptor modulators on a mouse model of fallopian tube epithelia, an ovarian cancer precursor | 8 | Mus musculus | https://www.ebi.ac.uk/arrayexpress/experiments/E-GEOD-67207/ |
| E-GEOD-67610 | Retinoids induce rapid dynamic changes in the non-coding RNAs and epigenetic profiles of murine Hox clusters | 15 | Mus musculus | https://www.ebi.ac.uk/arrayexpress/experiments/E-GEOD-67610/ |
| E-GEOD-67719 | Myc and SAGA Rewire an Alternative Splicing Network During Early Somatic Cell Reprogramming | 3 | Mus musculus | https://www.ebi.ac.uk/arrayexpress/experiments/E-GEOD-67719/ |
| E-GEOD-67828 | Profiling of soma and neurite transcriptomes | 7 | Mus musculus | https://www.ebi.ac.uk/arrayexpress/experiments/E-GEOD-67828/ |
| E-GEOD-67991 | Transcription profiling by high throughput sequencing of pancreatic islets isolated from free fatty acid receptor 3 knockout (Ffar3 KO) and wild type (Ffar3 WT) male mice | 2 | Mus musculus | https://www.ebi.ac.uk/arrayexpress/experiments/E-GEOD-67991/ |
| E-GEOD-68602 | Gene expression changes in control- and shSamd14-infected R1 fetal liver | 6 | Mus musculus | https://www.ebi.ac.uk/arrayexpress/experiments/E-GEOD-68602/ |
| E-GEOD-68617 | Trim33 binds and silences a class of young endogenous retroviruses in the mouse testis | 7 | Mus musculus | https://www.ebi.ac.uk/arrayexpress/experiments/E-GEOD-68617/ |
| E-GEOD-68715 | Global transcriptional analysis of mouse fibroblasts, chemically-induced neurons (neuron-like cells) from mouse fibroblasts and mouse primary cortical neurons by RNA- | 15 | Mus musculus | https://www.ebi.ac.uk/arrayexpress/experiments/E-GEOD-68715/ |
| E-GEOD-68902 | Direct Induction of Neurons from Various Cell Types by Chemical Defined Medium | 4 | Mus musculus | https://www.ebi.ac.uk/arrayexpress/experiments/E-GEOD-68902/ |
| E-GEOD-69018 | The impact of pregnancy experience on uterine gene expression and decidualization | 2 | Mus musculus | https://www.ebi.ac.uk/arrayexpress/experiments/E-GEOD-69018/ |
| E-GEOD-69180 | 1,25-Dihydroxyvitamin D3 Controls a Cohort of Vitamin D Receptor Target Genes in the Proximal Intestine That Is Enriched for Calcium Regulating Components | 3 | Mus musculus | https://www.ebi.ac.uk/arrayexpress/experiments/E-GEOD-69180/ |
| E-GEOD-69699 | Genome-wide translational analysis of the effect of acute endurance exercise on mouse gastrocnemius | 8 | Mus musculus | https://www.ebi.ac.uk/arrayexpress/experiments/E-GEOD-69699/ |
| E-GEOD-69913 | Age- and Obesity Induced Decline of Brown Fat Function as Consequence of Impaired miR-328 Dependent Silencing of Bace1 | 6 | Mus musculus | https://www.ebi.ac.uk/arrayexpress/experiments/E-GEOD-69913/ |
| E-GEOD-70016 | Spontaneous transformation of murine oviductal epithelial cells: A model system to investigate the onset of fallopian-derived tumors | 6 | Mus musculus | https://www.ebi.ac.uk/arrayexpress/experiments/E-GEOD-70016/ |
| E-GEOD-70123 | Adipose Rab10 Knockout Causes Insulin Resistance through Impaired Glucose Uptake | 6 | Mus musculus | https://www.ebi.ac.uk/arrayexpress/experiments/E-GEOD-70123/ |
| E-GEOD-70132 | BMP signaling differentially regulates distinct hematopoietic stem cell types | 4 | Mus musculus | https://www.ebi.ac.uk/arrayexpress/experiments/E-GEOD-70132/ |
| E-GEOD-70307 | global gene expression profiling experiments of HSC cultured in cysteine or valine deficient medium by RNA-sequencing analysis | 3 | Mus musculus | https://www.ebi.ac.uk/arrayexpress/experiments/E-GEOD-70307/ |
| E-GEOD-70608 | Single Cell Sequencing Identifies Key Epigenetic Regulators in Nuclear Transfer Mediated Reprogramming | 27 | Mus musculus | https://www.ebi.ac.uk/arrayexpress/experiments/E-GEOD-70608/ |
| E-GEOD-71145 | Cellular response to proteasome inhibition by Bortezomib | 6 | Mus musculus | https://www.ebi.ac.uk/arrayexpress/experiments/E-GEOD-71145/ |
| E-GEOD-71390 | The relationship between gene network structure and expression variation among individuals and species | 5 | Mus musculus | https://www.ebi.ac.uk/arrayexpress/experiments/E-GEOD-71390/ |
| E-GEOD-71405 | Analysis of transcriptomes during cardioac reprogramming by RNA-deep Sequencing | 5 | Mus musculus | https://www.ebi.ac.uk/arrayexpress/experiments/E-GEOD-71405/ |
| E-GEOD-71734 | RNA-seq data of intestinal epithelial cells | 16 | Mus musculus | https://www.ebi.ac.uk/arrayexpress/experiments/E-GEOD-71734/ |
| E-GEOD-72990 | Cells with Surface Expression of CD133highCD71low are Enriched for Colony-Forming Progenitor Cells in the Adult Murine Pancreas | 4 | Mus musculus | https://www.ebi.ac.uk/arrayexpress/experiments/E-GEOD-72990/ |
| E-GEOD-73049 | genes regulated by mitoflash | 5 | Mus musculus | https://www.ebi.ac.uk/arrayexpress/experiments/E-GEOD-73049/ |
| E-GEOD-73272 | RNA-Seq analysis of mouse group 2 innate lymphoid cells (ILC2s) cultured with IL-33, IFN-g, and IL-27 | 5 | Mus musculus | https://www.ebi.ac.uk/arrayexpress/experiments/E-GEOD-73272/ |
| E-GEOD-73611 | Combined loss of Tet1 and Tet2 promotes B-cell, but not myeloid malignancies in mice | 6 | Mus musculus | https://www.ebi.ac.uk/arrayexpress/experiments/E-GEOD-73611/ |
| E-GEOD-73631 | A XEN-like state bridges somatic cells to pluripotency during chemical reprogramming | 21 | Mus musculus | https://www.ebi.ac.uk/arrayexpress/experiments/E-GEOD-73631/ |
| E-GEOD-73711 | RNA sequencing of e15.5 pancreas from Wild Type, Blinc1-/- and Blinc+/- mice | 6 | Mus musculus | https://www.ebi.ac.uk/arrayexpress/experiments/E-GEOD-73711/ |
| E-GEOD-73861 | Development and Plasticity of Alveolar Type 1 Cells | 1 | Mus musculus | https://www.ebi.ac.uk/arrayexpress/experiments/E-GEOD-73861/ |
| E-GEOD-74064 | Therapeutic targeting of myeloid leukemias with spliceosomal mutations through modulation of splicing catalysis | 43 | Mus musculus | https://www.ebi.ac.uk/arrayexpress/experiments/E-GEOD-74064/ |
| E-GEOD-74371 | GATA-1 and heme regulate the erythroid cell transcriptome | 12 | Mus musculus | https://www.ebi.ac.uk/arrayexpress/experiments/E-GEOD-74371/ |
| E-GEOD-74855 | Transcriptional effects of I-PpoI induced DNA double strand breaks in mouse | 2 | Mus musculus | https://www.ebi.ac.uk/arrayexpress/experiments/E-GEOD-74855/ |
| E-GEOD-75103 | Population transcriptional profiling of in vitro polarized Th17 cells, either sorted for IL17A/GFP+ or unsorted | 7 | Mus musculus | https://www.ebi.ac.uk/arrayexpress/experiments/E-GEOD-75103/ |
| E-GEOD-75104 | Population transcriptional profiling of KO or WT cells,, differentiated in vitro for 48-96h towards Th17 cells | 30 | Mus musculus | https://www.ebi.ac.uk/arrayexpress/experiments/E-GEOD-75104/ |
| E-GEOD-75105 | Population transcriptional profiling of Th17 cells, isolated from CNS or LN at peak of disease in EAE | 6 | Mus musculus | https://www.ebi.ac.uk/arrayexpress/experiments/E-GEOD-75105/ |
| E-GEOD-75107 | Single-cell transcriptional profiling of Th17 cells, harvested at peak of disease in EAE from CNS [EAE-CNS-IL-17A/GFP+] | 165 | Mus musculus | https://www.ebi.ac.uk/arrayexpress/experiments/E-GEOD-75107/ |
| E-GEOD-75108 | Single-cell transcriptional profiling of Th17 cells, harvested at peak of disease in EAE from LN [EAE-LN-IL-17A/GFP+] | 136 | Mus musculus | https://www.ebi.ac.uk/arrayexpress/experiments/E-GEOD-75108/ |
| E-GEOD-75109 | Single-cell transcriptional profiling of Th17 cells, differentiated in vitro for 48h [IL1B IL6 IL23-48h-IL-17A/GFP+] | 139 | Mus musculus | https://www.ebi.ac.uk/arrayexpress/experiments/E-GEOD-75109/ |
| E-GEOD-75110 | Single-cell transcriptional profiling of Th17 cells, differentiated in vitro for 48h [TGFB1 IL6-48h] | 130 | Mus musculus | https://www.ebi.ac.uk/arrayexpress/experiments/E-GEOD-75110/ |
| E-GEOD-75111 | Single-cell transcriptional profiling of Th17 cells, differentiated in vitro for 48h [TGFB1 IL6-48h-IL-17A/GFP+] | 151 | Mus musculus | https://www.ebi.ac.uk/arrayexpress/experiments/E-GEOD-75111/ |
| E-GEOD-75522 | Pluripotent Stem Cells Induced from Mouse Neural Stem Cells and Small Intestinal Epithelial Cells by Small Molecule Compounds | 8 | Mus musculus | https://www.ebi.ac.uk/arrayexpress/experiments/E-GEOD-75522/ |
| E-GEOD-75544 | Sperm tsRNAs contribute to intergenerational inheritance of an acquired metabolic | 9 | Mus musculus | https://www.ebi.ac.uk/arrayexpress/experiments/E-GEOD-75544/ |
| E-GEOD-75615 | Distinct GAB2 signaling pathways are essential for myeloid and lymphoid transformation and leukemogenesis by BCR-ABL1 | 1 | Mus musculus | https://www.ebi.ac.uk/arrayexpress/experiments/E-GEOD-75615/ |
| E-GEOD-75788 | Loss of Asxl1 Alters Mesenchymal Stem Cell Fate through H3K4me3 | 3 | Mus musculus | https://www.ebi.ac.uk/arrayexpress/experiments/E-GEOD-75788/ |
| E-GEOD-76130 | Alternative routes to induced pluripotent stem cells revealed by reprogramming of neural lineage | 3 | Mus musculus | https://www.ebi.ac.uk/arrayexpress/experiments/E-GEOD-76130/ |
| E-GEOD-76143 | Zscan10 is dispensable for maintenance of pluripotency in mouse embryonic stem cells | 6 | Mus musculus | https://www.ebi.ac.uk/arrayexpress/experiments/E-GEOD-76143/ |
| E-GEOD-76288 | Impact of miRNA Levels, Target-Site Complementarity and Cooperativity on ceRNA-Regulated Gene Expression | 64 | Mus musculus | https://www.ebi.ac.uk/arrayexpress/experiments/E-GEOD-76288/ |
| E-GEOD-76356 | DICER controls macrophage polarization and tumor response to immunotherapy | 7 | Mus musculus | https://www.ebi.ac.uk/arrayexpress/experiments/E-GEOD-76356/ |
| E-GEOD-76412 | Hdac3 Interaction with p300 Histone Acetyltransferase Regulates the Oligodendrocyte and Astrocyte Lineage Fate Switch | 1 | Mus musculus | https://www.ebi.ac.uk/arrayexpress/experiments/E-GEOD-76412/ |
| E-GEOD-76579 | Single cell transcriptome analysis of mouse carotid body glomus cells | 8 | Mus musculus | https://www.ebi.ac.uk/arrayexpress/experiments/E-GEOD-76579/ |
| E-GEOD-76603 | Tumor suppressor role of Ezh2 in an NRASQ61K driven model of Early T-cell Precursor Acute Lymphoblastic Leukemia | 4 | Mus musculus | https://www.ebi.ac.uk/arrayexpress/experiments/E-GEOD-76603/ |
| E-GEOD-76606 | Asynchronous combinatorial action of four regulatory factors activates Bcl11b for T cell commitment | 6 | Mus musculus | https://www.ebi.ac.uk/arrayexpress/experiments/E-GEOD-76606/ |
| E-GEOD-76750 | The Histone Methyltransferases MLL1 and DOT1L Cooperate with Meningioma-1 to Induce AML | 11 | Mus musculus | https://www.ebi.ac.uk/arrayexpress/experiments/E-GEOD-76750/ |
| E-GEOD-76932 | Next generation sequencing comparing effects of vehicle, PTH, and YKL-05-093 in Ocy454 cells | 6 | Mus musculus | https://www.ebi.ac.uk/arrayexpress/experiments/E-GEOD-76932/ |
| E-GEOD-77744 | Essential role of the transcription factors E2A and E2-2 in germinal center B cell and plasma cell development | 13 | Mus musculus | https://www.ebi.ac.uk/arrayexpress/experiments/E-GEOD-77744/ |
| E-GEOD-77851 | Effects of Lin28a overexpression on gene expression changes during culturing neural stem cells (NSCs) derived from mouse embryonic ventral midbrain (VM) | 2 | Mus musculus | https://www.ebi.ac.uk/arrayexpress/experiments/E-GEOD-77851/ |
| E-GEOD-78902 | RNAseq analysis of the colon of intestine-specific adult Nkx2.2 mutant mice | 4 | Mus musculus | https://www.ebi.ac.uk/arrayexpress/experiments/E-GEOD-78902/ |
| E-GEOD-78965 | Differential gene expression analysis of follistain treated skeletal muscle | 9 | Mus musculus | https://www.ebi.ac.uk/arrayexpress/experiments/E-GEOD-78965/ |
| E-GEOD-78966 | Progenitor Cell Marker Aldehyde Dehydrogenase 1a3 Defines a Subset of Failing Pancreatic Beta Cells in Diabetic Mice | 13 | Mus musculus | https://www.ebi.ac.uk/arrayexpress/experiments/E-GEOD-78966/ |
| E-GEOD-79758 | Differential gene expression in intestinal mouse crypts after loss of KLF5 | 6 | Mus musculus | https://www.ebi.ac.uk/arrayexpress/experiments/E-GEOD-79758/ |
| E-GEOD-80089 | Gene expression profiling and genome-wide maps in PDGFB-induced glioma | 1 | Mus musculus | https://www.ebi.ac.uk/arrayexpress/experiments/E-GEOD-80089/ |
| E-GEOD-80669 | Defining AP4- and MYC-regulated gene expression programs in germinal center B cells during cyclic re-entry | 12 | Mus musculus | https://www.ebi.ac.uk/arrayexpress/experiments/E-GEOD-80669/ |
| E-GEOD-80678 | Gene expression signatures for primary tumor and tumor-derived exosomes | 4 | Mus musculus | https://www.ebi.ac.uk/arrayexpress/experiments/E-GEOD-80678/ |
| E-GEOD-80804 | A Pro-Asthmatic IL-4 Receptor Allele Promotes Airway Inflammation by Programming TH17 Cell-Like Regulatory T cells | 7 | Mus musculus | https://www.ebi.ac.uk/arrayexpress/experiments/E-GEOD-80804/ |
| E-GEOD-81045 | Zfp281 Coordinates Opposite Functions of Tet1 and Tet2 for Alternative Pluripotent | 14 | Mus musculus | https://www.ebi.ac.uk/arrayexpress/experiments/E-GEOD-81045/ |
| E-GEOD-82471 | RNA-seq from heart (ENCSR150CUE) | 2 | Mus musculus | https://www.ebi.ac.uk/arrayexpress/experiments/E-GEOD-82471/ |
| E-GEOD-82852 | RNA-seq from forebrain (ENCSR647QBV) | 2 | Mus musculus | https://www.ebi.ac.uk/arrayexpress/experiments/E-GEOD-82852/ |
| E-GEOD-82853 | RNA-seq from liver (ENCSR648YEP) | 2 | Mus musculus | https://www.ebi.ac.uk/arrayexpress/experiments/E-GEOD-82853/ |
| E-GEOD-82919 | RNA-seq from limb (ENCSR750YSX) | 2 | Mus musculus | https://www.ebi.ac.uk/arrayexpress/experiments/E-GEOD-82919/ |
| E-GEOD-82976 | RNA-seq from embryonic facial prominence (ENCSR851HEC) | 2 | Mus musculus | https://www.ebi.ac.uk/arrayexpress/experiments/E-GEOD-82976/ |
| E-GEOD-83966 | Sox2 suppresses gastric tumorigenesis in mice | 6 | Mus musculus | https://www.ebi.ac.uk/arrayexpress/experiments/E-GEOD-83966/ |
| E-GEOD-84987 | Molecular role of the PAX5-ETV6 oncoprotein in promoting B cell acute lymphoblastic leukemia | 24 | Mus musculus | https://www.ebi.ac.uk/arrayexpress/experiments/E-GEOD-84987/ |
| E-GEOD-84989 | Genome-wide analysis of the transcriptional and alternative splicing landscape in intestinal organoids undergoing nutrient starvation or ER stress | 12 | Mus musculus | https://www.ebi.ac.uk/arrayexpress/experiments/E-GEOD-84989/ |
| E-MTAB-2836 | RNA-seq of coding RNA from tissue samples of 122 human individuals representing 32 different tissues | 37 | Homo sapiens | https://www.ebi.ac.uk/arrayexpress/experiments/E-MTAB-2836/ |
| E-GEOD-43717 | Transcription profiling by high throughput sequencing of purified Sertoli cells, spermatogonia, spermatocytes, spermatids and spermatozoa from mice | 5 | Mus musculus | https://www.ebi.ac.uk/arrayexpress/experiments/E-GEOD-43717/ |
