## Supplementary table 3 for "FertilityOnline, a straight pipeline for functional gene annotation and disease mutation discovery, identifies novel infertility causative mutations in *SYCE1* and *STAG3*"

**Table S3. Curated testicular scRNA-seq datasets**

| Accession | Species | Library preparation method | Year | PMID |
| --- | --- | --- | --- | --- |
| E-MTAB-6946 | mouse | 10X Chromium | 2019 | 30890697 |
| GSE107644 | mouse | Smart-seq2 | 2018 | 30061742 |
| GSE120508 | human | 10X Chromium | 2018 | 30315278 |
| GSE106487 | human | Smart-seq2 | 2018 | 30174296 |
