## Supplementary table 4 for "FertilityOnline, a straight pipeline for functional gene annotation and disease mutation discovery, identifies novel infertility causative mutations in *SYCE1* and *STAG3*"

**Table S4. The performance of variant annotation using FertilityOnline**

| Run | Variants | Total (s) | Step1 (s) | Step2 (s) | Step3 (s) | Step4 (s) | Step5 (s) | Step6 (s) |
|---|---|---|---|---|---|---|---|---|
| 1 | 175163 | 213.1 | 4.8 | 22.2 | 73.5 | 88.7 | 5.3 | 18.6 |
| 2 | 183762 | 216.6 | 5 | 22.5 | 74.7 | 90 | 5.4 | 19 |
| 3 | 315843 | 389.2 | 5.5 | 32.4 | 158.1 | 159.2 | 6.3 | 27.7 |
| 4 | 421137 | 375.5 | 6 | 40.7 | 107.8 | 180.1 | 6.9 | 34 |
| 5 | 307059 | 328.7 | 5.5 | 31.9 | 100.1 | 158.6 | 6.2 | 26.4 |
| 6 | 370316 | 373.4 | 5.8 | 35.4 | 125.8 | 169.1 | 6.8 | 30.5 |
| 7 | 368266 | 369.5 | 5.8 | 34.7 | 120 | 172.7 | 6.7 | 29.6 |
| 8 | 168171 | 258.6 | 4.8 | 21.4 | 112.3 | 95.9 | 5.3 | 18.9 |
| 9 | 132890 | 238.7 | 4.6 | 19.4 | 107.5 | 85.9 | 5.1 | 16.2 |
| 10 | 173927 | 246.5 | 4.9 | 21.8 | 103.7 | 92.1 | 5.4 | 18.6 |
| Average | 261653.4 | 300.98 | 5.27 | 28.24 | 108.35 | 129.23 | 5.94 | 23.95 |

**Notes:** Step1: Parse input and detect the format of input; Step2: Annotation of variant consquence; Step3: Dleteriousness prediction; Step4: Annotaion of MAF; Step5: Annotation of gene function information; Spte6: Integration of all results.
