## Supplementary table 5 for "FertilityOnline, a straight pipeline for functional gene annotation and disease mutation discovery, identifies novel infertility causative mutations in *SYCE1* and *STAG3*"

**Table S5. Statistical results of reproted functional genes based on species**

| **Species** | **Gene counts** | **Percentage** |
|---|---|---|
| *Mus musculus* | 1028 | 61.59 |
| *Homo sapiens* | 264 | 15.82 |
| *Drosophila melanogaster* | 167 | 10.07 |
| *Rattus norvegicus* | 44 | 2.64 |
| *Caenorhabditis elegans* | 31 | 1.86 |
| Others | 135 | 8.03 |
