## Supplementary table 6 for "FertilityOnline, a straight pipeline for functional gene annotation and disease mutation discovery, identifies novel infertility causative mutations in *SYCE1* and *STAG3*"

**Table S6. Statistical results of reported functional genes based on functional stages**

| Functional stage | | Gene counts | | Percentage (%) | |
|---|---|---|---|---|---|
| Premeiotic | Premeiotic only | 146 | 579 | 9.07 | 35.96 |
| | Premeioti / Meiotic | 112 | | 6.96 | |
| | Premeiotic / Postmeiotic | 21 | | 1.30 | |
| | Premeiotic / Meiotic / Postmeiotic | 300 | | 18.63 | |
| Meiotic | Meiotic only | 357 | 994 | 22.17 | 61.74 |
| | Premeiotic / Meiotic | 112 | | 6.96 | |
| | Meiotic / Postmeiotic | 225 | | 13.98 | |
| | Premeiotic / Meiotic / Postmeiotic | 300 | | 18.63 | |
| Postmeiotic | Postmeiotic only | 455 | 1001 | 28.26 | 62.17 |
| | Premeiotic / Postmeiotic | 21 | | 1.30 | |
| | Meiotic / Postmeiotic | 225 | | 13.98 | |
| | Premeiotic / Meiotic / Postmeiotic | 300 | | 18.63 | |
| Unclear | / | 63 | 63 | 3.77 | 3.77 |
