## Supplementary table 7 for "FertilityOnline, a straight pipeline for functional gene annotation and disease mutation discovery, identifies novel infertility causative mutations in *SYCE1* and *STAG3*"

**Table S7. Statistical results of the reported functional genes based on cell types**

| Functional cell type | Gene counts | Percentage (%) |
|---|---|---|
| Spermatocyte | 849 | 52.73 |
| Spermatid | 816 | 50.68 |
| Spermatogonia | 384 | 23.85 |
| Sertoli cell | 268 | 16.65 |
| Elongated spermatid | 149 | 9.25 |
| Leydig cell | 114 | 7.08 |
| Spermatogonial stem cell | 62 | 3.85 |
| Sperm | 29 | 1.80 |
| Round spermatid | 8 | 0.50 |
| Gonocyte | 4 | 0.25 |
| Elongating spermatid | 4 | 0.25 |
| Germline stem cell | 1 | 0.06 |
